# Chemically induced USP21-KLHL12 association recruits USP21 to COPII assembly

**DOI:** 10.64898/2026.09.13.750941

**Authors:** Victoria Vu, Michael Kanaris, Esther Wolf, Sabrina Keller, Qi Liu, Conrad Veranso Simoben, Shabbir Ahmad, Viviane Reber, Magdalena M Szewczyk, Peter Loppnau, Matthew ER Maitland, Ella C Adamson, Paola Picotti, Derek Wilson, Cheryl Arrowsmith, Levon Halabelian, Matthieu Schapira, Matthias Gstaiger, Rachel J. Harding, Dalia Barsyte-Lovejoy

## Abstract

Ubiquitin-specific protease 21 (USP21) is a deubiquitinase implicated in various biological processes and disease states, including cancer. To better understand USP21 protein interaction networks, we used biotin identification proximity labeling, and uncovered USP21 interactions with the cytoskeleton, endosomal, and transcriptional regulators. Notably, a subset of these interactions was modulated by the USP21-selective inhibitor BAY-805. In cells, BAY-805 promoted proximity between USP21 and Kelch-like protein KLHL12, a component of the Cullin3 ligase complex. Structural modeling using AlphaFold-3, together with hydrogen–deuterium exchange mass spectrometry (HDX-MS), identified an orthosteric BAY-805 binding site within USP21. Binding of BAY-805 functionally competes with ubiquitin at USP21, thereby enabling USP21 interaction with KLHL12. This BAY-805 induced proximity elicited relocalization of USP21 to KLHL12 containing coat protein complex II (COPII) vesicles. Induced USP21–KLHL12 association led to enlargement of COPII vesicles and impaired transport of large cargoes such as collagen. Overall, our findings reveal a novel chemically induced proximity mechanism in which the monovalent ligand BAY-805 outcompetes ubiquitin from USP21, thereby recruiting USP21 to KLHL12 associated protein complexes.

## INTRODUCTION

The ubiquitin system regulates a wide range of cellular processes through the reversible conjugation of ubiquitin to target proteins. Conjugation of ubiquitin is mediated by an E1–E2–E3 ligase cascade, whereas its removal is catalyzed by deubiquitinases (DUBs). This dynamic regulation underlies numerous cellular functions, including proteasomal protein turnover, rapid inflammatory responses, DNA repair, membrane trafficking, and diverse signaling pathways. ^1, 2^

Ubiquitin specific protease 21 (USP21) is a DUB that has been reported to deubiquitinate a broad spectrum of substrates ^3^, such as histone H2A lysine 119 monoubiquitination (H2AK119ub) ^4–6^, Nanog ^7^, and mitogen activated protein kinase kinase 2 ^8^, among others. Through these activities, USP21 has been implicated in regulating chromatin organization, stem cell maintenance, and growth factor signaling. USP21 has also been linked to inflammatory and antiviral responses by deubiquitinating absent in melanoma 2 ^9^, stimulator of interferon genes (STING) ^10^, and receptor interacting protein 1 (RIP1) ^11^, thereby influencing inflammasome activation and interferon-mediated signaling pathways. At the subcellular level, USP21 displays prominent association with the cytoskeleton, including binding to microtubules and centrosomal components. In this context, USP21 also regulates GLI1 transcription factor signaling by recruiting and stabilizing GLI1 at the centrosome, where it promotes protein kinase A–dependent phosphorylation and activation of GLI1. ^3, 12, 13^

Selective inhibitors of deubiquitinases (DUBs) have enabled therapeutic targeting of this enzyme class and substantially advanced our understanding of DUB function ^14^. Several small molecule inhibitors of USP21 have been reported to date. (reviewed ^3, 15^) Recently, we described BAY-805, the first selective chemical probe targeting USP21, which exhibits nanomolar potency both in vitro and in cells.^16^

To further define the cellular consequences of USP21 inhibition by BAY-805, we mapped the USP21 interaction landscape in its native cellular context using biotin identification (BioID), a proximity-based labeling approach. This analysis revealed an extensive USP21 proximal interaction network enriched for microtubule-associated and cytoskeletal processes. Notably, however, BAY-805 treatment altered only a small subset of USP21 interactions. Among these, BAY-805 selectively induced proximity between USP21 and the Kelch-like protein KLHL12, a substrate receptor of the Cullin 3 (CUL3) E3 ligase complex. Mechanistically, we defined an orthosteric binding mode for BAY-805 within USP21, in which compound engagement precludes ubiquitin binding. This competitive displacement enables the interaction between USP21 and KLHL12, establishing a chemically induced proximity driven by a monovalent ligand. Functionally, BAY-805 dependent interaction promotes colocalization of USP21 with KLHL12 containing coat protein complex II (COPII) assemblies, leading to enlargement of COPII vesicles and disruption of transport of large secretory cargoes.

## RESULTS

### Identification of USP21-associated protein networks

To elucidate the in situ USP21 interactome, we performed biotin identification (BioID), a proximity-dependent biotinylation approach, followed by protein identification using bottom-up tandem mass spectrometry (MS/MS). Cells with inducible full length USP21 fused to miniTurbo at either the N- or C- termini or miniTurbo vector control were generated (**Figure 1A, and S1**). Biotinylated proteins were affinity purified, analyzed by MS/MS and using stringent filtering criteria, we identified 466 proteins that were significantly enriched in both N and C terminal USP21–miniTurbo datasets compared with miniTurbo alone. High confidence interactors were defined as proteins detected in less than 20% of the CRAPome BioID background database ^17^ and exhibiting significant enrichment (log2 fold change ≥ 2, P ≤ 0.01) (**Figure 1B**). Gene Ontology (GO) term analysis revealed enrichment for microtubule and centrosome-associated components, consistent with previous reports of USP21 localization and function ^12^ (**Figure 1C**). A substantial fraction of USP21 proximal interactors comprised proteins involved in the microtubule organizing center, microtubules, centrosomes, endosomes, processing bodies (P bodies), and the homologous to augmin subunits (HAUS) complex (**Figure 1D**), highlighting proximity of USP21 to cytoskeletal and membrane-associated structures and supporting previous USP21 associations with centrosome and cytoskeleton.^12^

**Figure 1.**
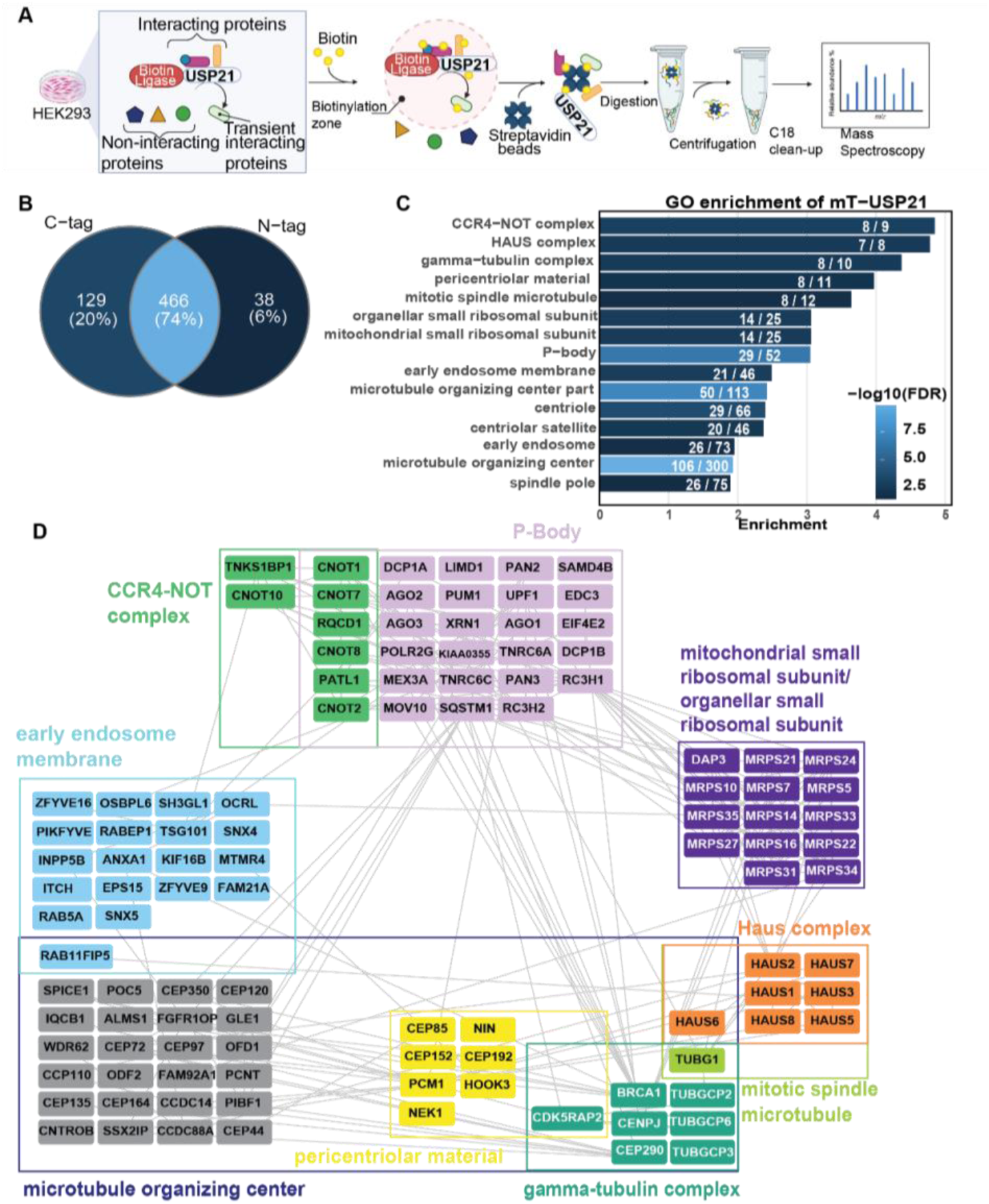
BioID identifies USP21 associated proteins are involved in a broad range of cellular pathways. (A) Scheme for USP21 proximity profiling using miniTurboID-MS. Made with Biorender. (B) Summary of USP21 associated proteins as Venn diagram for USP21 tagged with miniTurbo at the C- or N-terminus. Only proteins detected in less than 20% of the Crapome BioID background database and significantly enriched (log2FC ≥ 2, adjusted p-value ≤ 0.01) compared to miniTurbo control cells were considered. (C) The top 15 gene ontology categories of significantly enriched USP21 associated proteins (see Suppl. Table 1). Numbers inside the bars indicate the enrichment (odds ratio). (D) Interaction network of proteins matching the top 10 enriched GO terms from (C) of the USP21 proximity-based on BioGrid interaction data.

### USP21 chemical probe BAY-805, induces an association between USP21 and KLHL12

BAY-805 is a selective catalytic inhibitor of USP21 with nanomolar potency, although its precise binding site was unclear.^16^ To determine whether USP21 inhibition by BAY-805 alters its protein interaction landscape, we compared USP21 proximity labeling profiles following treatment with BAY-805, vehicle control (DMSO), or an inactive negative control compound (BAY-805NC). Across both comparison groups (BAY-805 vs DMSO and BAY-805 vs BAY-805NC), BAY-805 treatment resulted in a limited number of significantly increased or decreased USP21 interactions (**Figure 2A–B and S2A–B**).

**Figure 2.**
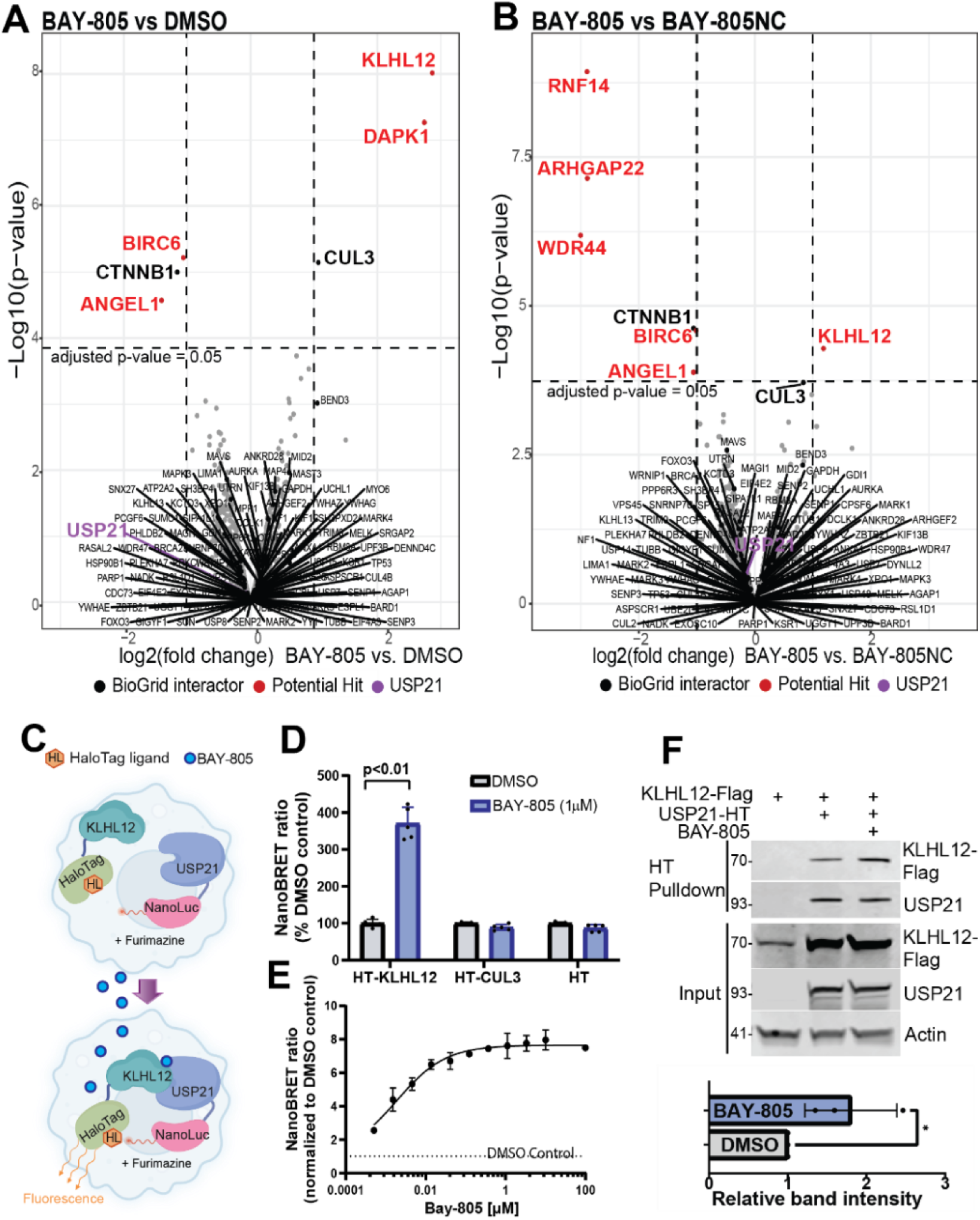
USP21 chemical probe BAY-805, induces an association between USP21 and KLHL12. (A) USP21 proximity profiling of HEK293 cells expressing C-tagged miniTurbo-USP21. Volcano plot displaying the log2 fold changes (x-axis) and the log10 p-values (y-axis) for proteins of the USP21 proximity in the presence of BAY-805 (1µM) compared to DMSO. Significance cut offs are indicated by dashed lines (log2FC ≥ |1| and adjusted p-value ≤ 0.05). (B) Volcano plot as in (A) displaying the proteins of the USP21 proximity in the presence of BAY-805 (1µM) compared to negative control BAY-805NC (1µM). Significance cut offs are indicated by dashed lines (log2FC ≥ |1| and adjusted p-value ≤ 0.05). (C) Schematic of NanoBRET experimental design to validate the interaction of USP21 with KLHL12 upon addition of BAY-805. NanoLuciferase (NanoLuc) tagged USP21 generates luminescence signal that when in proximity with HaloTag tagged KLHL12 transfers energy to the HaloTag ligand resulting in its fluorescence. (D) BAY-805 induces an increase in interaction between USP21 and KLHL12 in NanoBRET assay. HEK293T cells were transfected with NanoLuc (NL)-tagged USP21 and HaloTag (HT)-tagged KLHL12, CUL3 or HT alone and treated with 1 µM BAY-805 for 2 h. The nanoBRET ratio was measured by dividing intensity of acceptor fluorescent signal by donor bioluminescence signal (n=6, two independent experiments). Statistical significance was determined with unpaired Student *t*-test (two-tailed). (E). BAY-805 increases the NanoBRET ratio of USP21-NL and HT-KLHL12 in a dose-dependent manner. The graph represents non-linear fit of the NanoBRET ratio of cells treated with BAY-805 normalized to DMSO control: *n* = 3, biological replicates, EC_50_ = 2nM (F). BAY-805 facilitates USP21 and KLHL12 association in cells. HEK293 cells were transfected with tagged proteins, treated with BAY-805 and lysates subjected to immunoprecipitation blotting followed by western blotting. 3 biological replicates, unpaired Student t-test (two-tailed), p value <0.05.

BAY-805 treatment decreased the association of USP21 with the known interactor β-catenin (CTNNB1) ^18^, as well as with BIRC6 and ANGEL1 (**Figure 2A–B**). Unexpectedly, however, KLHL12 emerged as a high-confidence interactor significantly enriched in both BAY-805 comparison groups (**Figure 2A–B**), suggesting that BAY-805 induces proximity between USP21 and KLHL12.

The BAY-805 dependent USP21–KLHL12 interaction was independently confirmed using a cellular *in situ* NanoLuciferase (NL) bioluminescence resonance energy transfer (NanoBRET) assay ^19^, where USP21 fused to NL served as the energy donor, while KLHL12 was fused to a HaloTag (HT) protein binding a 618 nm fluorophore and acted as the BRET acceptor (**Figure 2C–D**). BAY-805 induced a robust increase in USP21–KLHL12 proximity across all combinations of N and C terminal protein tagging. Notably, BioID analysis did not reveal a significant enrichment of KLHL12 with N-terminally miniTurbo tagged USP21, likely reflecting unfavorable spatial orientation for efficient biotinylation rather than absence of interaction (**Figure S2A–C**).

We next used NanoBRET to assess whether BAY-805 affected established USP21 interactions with CTNNB1, ^18^ KCTD6, ^13^ as well as newly identified candidates. Although CTNNB1, KCTD6, DAPK1, and ANGEL1 all interacted with USP21, BAY-805 did not significantly alter these associations. In contrast, a modest increase in USP21 interaction with DAPK1 was observed upon BAY-805 treatment (**Figure S2D**).

KLHL12 functions as a substrate adaptor within a Cullin RING ligase (CRL) complex containing CUL3. ^20, 21^ Given that CUL3 was also detected in the USP21 BioID dataset (**Figure 2A**), we tested whether BAY-805 promotes interaction between USP21 and CUL3. NanoBRET analysis revealed that BAY-805 selectively induced USP21 association with KLHL12 but not CUL3 (**Figure 2D**), indicating potentially direct interaction with the substrate adaptor. Dose–response analysis demonstrated that BAY-805 induced USP21–KLHL12 complex formation with an EC₅₀ of approximately 2 nM, without evidence of a “hook effect” at concentrations up to 100 µM (**Figure 2E**). This BAY-805 dependent USP21–KLHL12 association was confirmed by USP21 pull-down from cell lysates, providing orthogonal biochemical validation (**Figure 2F**).

### USP21 structural determinants of BAY-805 binding

Protein sequence alignment of USP21 with its closest USP family homologs revealed a unique insertion located between the thumb α5 helix and the fingers α6 helix, corresponding to the region between conserved USP boxes 2 and 3 (**Figure 3A**).^22^ This insertion is largely This insertion is unresolved in experimental structures solved to date and is predicted to be disordered in AF3 models. Pronounced insertions are present in USP21, USP10, and USP2, whereas this region is absent in USP7. Attempts to co-crystallize BAY-805 with USP21 were unsuccessful, so we turned to limited proteolysis coupled to mass spectrometry (LiP MS) to identify USP21 regions whose protease sensitivity changes upon ligand binding (**Figure 3B**). Analysis of peptide intensities across increasing concentrations of BAY-805 identified a single peptide spanning residues 323–338 that exhibited a significant, dose-dependent decrease in abundance. Quantitative fitting of the response curve (correlation coefficient = 0.9) yielded a predicted EC₅₀ of 63 nM. No change in proteolytic susceptibility was observed upon treatment with the negative control compound, indicating specificity. This differential proteolysis suggests that the 323–338 region either directly contributes to BAY-805 binding or undergoes ligand-induced conformational changes. Consistent with this interpretation, sequence comparison places this peptide within the disordered insertion between conserved USP boxes 2 and 3 (**Figure 3A**).

**Fig 3.**
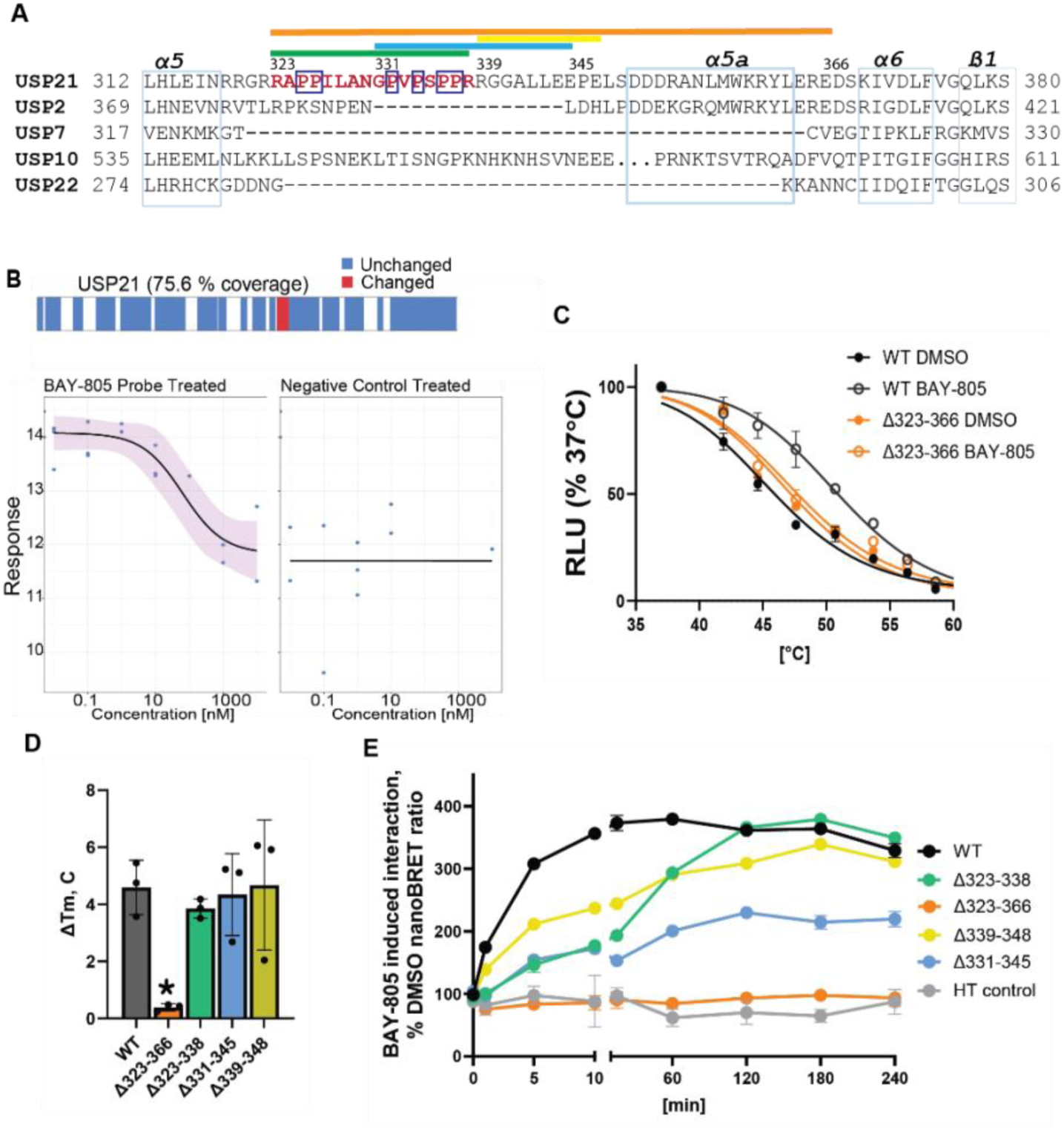
Extended region of USP21 between thumb and fingers subdomains is required for binding of BAY-805 and KLHL12. (A) Alignment of USPs most closely related to USP21. LiP-MS detected peptide is indicated in red. Light blue boxes show conserved thumb helix α5, and fingers α6, β1 and non conserved α5a regions. Dark blue boxes indicate prolines within LiP-MS detected peptide that potentially bind to KLHL12. The color lines at the top indicate regions deleted in mutational analysis (Fig 3E). (B) LiP-MS peptide coverage of USP21 protein with blue labels as peptides detected but not changed while peptide in red (RAPPILANGPVPSPPR) was differentially detected upon BAY-805 treatment. Bottom graph indicates dose response of the peptide detection upon treatment with BAY-805 or negative control compound. (C) NanoLuciferase cellular Thermal Shift Assay (NalTSA) indicates that BAY-805 (1 μM) stabilizes WT USP21 but not 323-366 region deleted USP21. (D) NalTSA ΔT_m_ summary indicates that only 323-366 region deletion abrogates BAY-805-mediated stabilization. Star indicates p value <0.05. (E) Deletions within the 323-366 region of USP21 reduce the BAY-805 induced USP21-KLHL12 association. Experiments performed as in B. The results are Mean+/-SD (n=3). Note that the deletions are color-coded as in panel A.

We next investigated whether the same LiP MS identified region mediates BAY-805 binding to USP21 and association with KLHL12. Based on the LiP MS findings and prior reports highlighting the importance of proline-rich motifs in KLHL12 substrate recognition, ^23–25^ we generated a series of USP21 deletion constructs encompassing the LiP MS peptide and neighboring proline-rich regions. We also included a larger 323–366 deletion that mimics the absence of this extended insertion, as observed in USP7 and USP22 (**Figure 3A**).

To determine whether this region is required for cellular target engagement by BAY-805, we employed a NanoLuciferase based cellular thermal shift assay (NalTSA).^26^ BAY-805 induced a robust increase in the melting temperature (Tₘ) of wild type USP21, whereas deletion of residues 323–366 completely abolished BAY-805 dependent thermal stabilization (**Figure 3C–D**). Importantly, these deletions did not affect USP21 expression levels or Tₘ (**Figure S3A–B**), indicating that loss of stabilization was not due to protein instability.

We monitored USP21–KLHL12 association kinetics over a 4-hour time course. BAY-805 rapidly induced USP21–KLHL12 interaction, with the NanoBRET signal reaching half-maximal levels within five minutes, consistent with efficient compound uptake and rapid complex formation (**Figure 3E**). Deletion of the entire insertion region (Δ323–366), including the LiP MS identified peptide, completely abolished BAY-805 induced interaction with KLHL12 (**Figure 3E**). In contrast, deletion of only the LiP MS peptide (Δ323–338) did not prevent BAY-805 dependent KLHL12 binding, consistent with NalTSA results (**Figure 3C–D**).

We also assessed whether BAY-805 can induce KLHL12 association with related USPs. No such induced association was observed with USP2, USP10, or USP22 (**Figure S3C**). Furthermore, although LiP MS was performed using the isolated catalytic domain of USP21 (residues 209–563), whereas NalTSA and NanoBRET experiments used full-length protein, we confirmed that the USP21 catalytic domain alone is sufficient to support BAY-805 induced KLHL12 interaction (**Figure S3D**). Collectively, these data identify the USP21 323–366 region as a critical determinant for BAY-805 mediated USP21–KLHL12 interaction; however, proline-rich region identified by LiP MS was not critical for USP21-KLHL12 association elicited by BAY-805.

### BAY-805 computationally co-folds with USP21

To gain additional structural insight into BAY-805 binding, we generated cofolded models of the USP21–BAY-805 complex using AlphaFold 3 (AF3), Boltz 2, and Chai 1. Among these approaches, only AF3 produced a high confidence ligand bound model (ligand pLDDT = 94) (**Figure 4A and Fig S4A-B**). In this model, BAY-805 occupies the orthosteric S1 binding site of USP21, spanning two adjacent cavities centered around residues Q377 and C398, separated by V396 (**Figure 4A–B**). The model predicts that the cyano group of the benzonitrile moiety of BAY-805 forms a hydrogen bond with the side chain carboxamide of Q377, potentially anchoring the ligand within the pocket (**Figure 4C**). In addition, the thiadiazole ring is positioned to engage in weak cation–π interactions with R365, suggesting an electrostatic contribution to ligand stabilization. The thiazole amine is predicted to form a hydrogen bond with the carboxylate of E366, while the phenolic hydroxyl group of Y362 is predicted to hydrogen bond with the distal carboxamide of the ligand. Together, these interactions may orient the cyclohexyl substituent of BAY-805 deeply within an adjacent hydrophobic cavity (**Figure 4B–C**).

**Figure 4.**
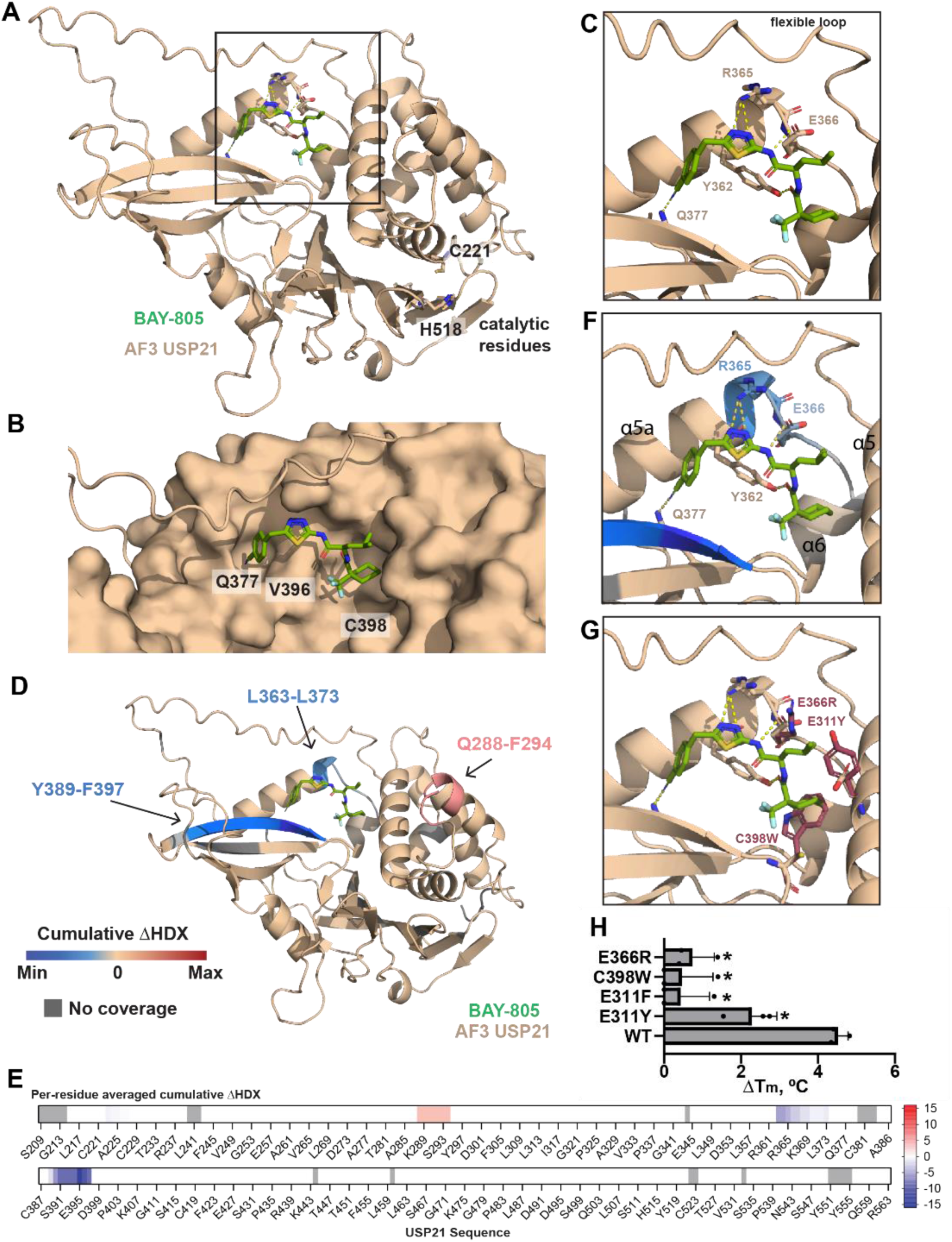
BAY-805 engages orthosteric S1 site of USP21. (A). AlphaFold 3 model of USP21-BAY-805 binding. USP21 AF3 structure (tan) with docked BAY-805 (green) (B). Surface representation of BAY-805 binding pocket with C398 and Q377 cavities denoted and flexible loop shown as a line. (C) Critical USP21 residues (sticks) predicted to form hydrogen bonds (yellow dashes) with BAY-805. (D). Visualization of ΔHDX-MS heatmap. ΔHDX-MS was plotted on AF3 model with the docked BAY-805 molecule (green) overlayed. Blue, tan, and grey indicate regions of decreased ΔHDX, no change, and no coverage respectively. (E). Per-residue averaged cumulative ΔHDX-MS. In the presence of BAY-805, decreases in HDX are shown in blue, increases in red, and no change is shown in white. Areas lacking peptide coverage are grey. (F). Closeup of ΔHDX and key residues identified by in silico prediction. The conserved α5 and α6 helices and α5a helix in flexible insertion region are indicated. (G). Point mutants (red sticks) of USP21 predicted to abrogate BAY-805 binding. (H) NalTSA validation of predicted residues (in G). Data presented as a summary of T_m_ shifts, N=3. Stars indicate significance (p<0.05) when compared to WT, one-way ANOVA, Dunnetts posthoc test.

Although the overall protein fold and the orthosteric S1 binding site were predicted with high confidence (average protein pLDDT = 87), the conformation of the flexible LiP MS identified peptide region (residues 323–338) was modeled with substantially lower confidence (pLDDT ≈ 60), consistent with its intrinsically disordered nature (**Figure S4C**). This is consistent with deletion of residues 323–338 not leading to impaired BAY-805 binding in cellular target engagement assays (**Figure 3D**), indicating that this flexible loop is unlikely to contribute directly to ligand binding.

### HDX-MS demonstrates BAY-805 binding to the orthosteric site of USP21, consistent with the AF3 model

Next, we performed differential hydrogen-deuterium exchange mass spectrometry (ΔHDX-MS) to test how the conformational dynamics of the catalytic domain of USP21 (residues 209-563) were impacted by BAY-805 (**Figure 4D-E**). The cumulative ΔHDX, illustrated in **Figure S4D-E** revealed that relative to USP21 (residues 209-563) alone, incubation with BAY-805 decreased deuterium uptake in peptides spanning residues 363-373 and 389-397. Per-residue analysis of each peptide region, enabled by redundant peptides, revealed that the highest magnitude ΔHDX occurred at amino acids L363, E364, E395, and V396 (**Figure 4D-F**). These results are consistent with the region identified in the docked BAY-805 AF3 structure (**Figure 4A-C**) and lie within or proximal to the broader 323-366 region identified as critical for BAY-805 binding (**Figure 3**). Interestingly, a short region spanning Q288-F294 exhibited increased ΔHDX, though the cumulative signals exceeding error at this timescale were subtle (+∼0.4 %). This region is localized and spatially distinct from the BAY-805 pocket, suggesting increased flexibility upon compound binding.

To guide further experimental confirmation of the predicted ligand binding mode by site directed mutagenesis, sidechains lining the allosteric pocket were mutated *in silico*, followed by molecular dynamics simulations to evaluate the stability of the mutated protein with and without BAY-805. Mutations C398W, E311Y, E311F, and E366R were predicted to destabilize ligand binding without affecting the apo protein structure (**Figure 4G**). We next tested these predictions experimentally by assessing BAY-805 induced thermal stabilization of USP21 mutants. Mutations E311Y, E311F predicted to sterically hinder the isobutyl group of BAY-805, reduced, and C398W (steric clash) and E366R (charge swap) abolished the compound induced thermal stabilization (**Figure 4H and Figure S4F**). *In silico* modeling predicted that these mutations would not disrupt protein structure, which was the case for E311Y/F and E366R, but the C398W mutation resulted in markedly reduced protein expression levels (**Figure S4G, H**). Together, the orthosteric binding mode is supported by HDX-MS decreases at residues 363–373 and 389–397 and by loss of stabilization in E366R and C398W mutants, consistent with occupancy of the S1 site.

### KLHL12 KELCH domain dimerization is required for the BAY-805-induced interaction with USP21

To further define the mechanism by which BAY-805 induces recruitment of USP21 to KLHL12, we investigated which domains of KLHL12 are required for this interaction. In KLHL12, the BTB domain mediates homodimerization of Kelch proteins and association with Cullin RING ligase complexes, whereas the BACK domain is proposed to contribute to complex assembly and substrate positioning. The C-terminal Kelch domain is responsible for substrate recognition (**Figure 5A**).^23, 27, 28^

**Fig 5.**
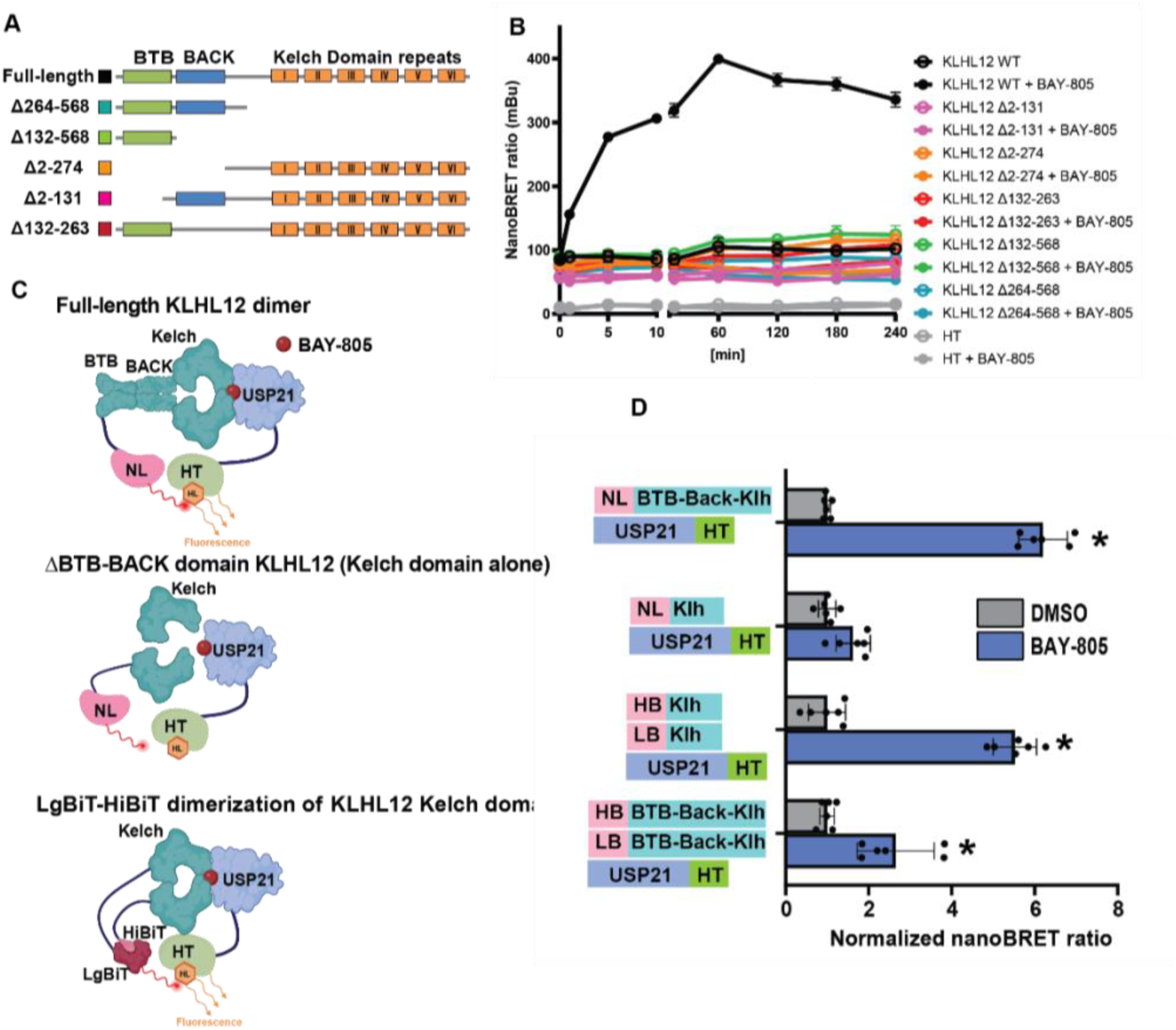
KLHL12 KELCH domain dimerization is required for BAY-805 induced association with USP21. (A) Schematic of KLHL12 mutations used in the USP21-KLHL12 interaction nanoBRET assays. (B) All deletions in KLHL12 abolish the BAY-805 induced KLHL12-USP21 association. NanoBRET testing interactions between USP21 and different KLHL12 mutants was performed in HEK293 cells as in Fig 4B. (C) Schematic of LgBiT-HiBiT induced dimerization of KLHL12. Panels show the dimerization of full length KLHL12 when used in NanoLuc (NL) KHL12 and USP21-HT NanoBRET assay. When BTB-BACK domain of KLHL12 is deleted, the Kelch domains no longer form dimers. However, when HiBiT-tagged KLHL12 Kelch domain is co-expressed with LgBiT-tagged KLHL12 Kelch domain, the complementation of HiBiT and LgBiT into functional NanoLuc enzyme promotes KLHL12 Kelch domain dimerization and the BAY-805 dependent interaction with USP21-HT. (D) HiBiT (HB) and LgBiT (LB) complementation induces Kelch domain dimerization, which results in KLHL12 and USP21 association upon BAY-805 treatment. HEK293T cells were transfected with indicated construct pairs for 24 h and treated with BAY-805 (1 µM) for 1 h. The full length KLHL12 was used as control. The results are mean of 2 independent triplicate experiments, stars indicated significant p<0.05 signal increase from respective control (one-way ANOVA, Dunnetts posthoc test).

Because the Kelch domain mediates direct substrate recognition, we initially hypothesized that deletion of this domain alone would abolish USP21 binding. However, NanoBRET analysis assessing interactions between wild type USP21 and the KLHL12 deletion variants revealed that BAY-805 induced USP21–KLHL12 interaction was lost not only upon removal of the Kelch domain, but also upon deletion of either the BTB or BACK domains (**Figure 5B**). Members of the KLHL protein family are known to form homodimers, which facilitate the coordinated recruitment of substrates and co-adaptors.^29^ To test whether KLHL12 dimerization is essential for BAY-805 driven USP21 recruitment, we employed a split NanoLuciferase system in which luminescence is reconstituted through high affinity binding between LgBiT and HiBiT tags.^30^ By tagging the isolated Kelch domain of KLHL12 through a linker to LgBiT and HiBiT, we reconstituted Kelch–Kelch dimerization independently of the BTB BACK domains (**Figure 5C**).

Strikingly, upon BAY-805 treatment, dimerized Kelch domains interacted with USP21 to a similar extent as full length NanoLuciferase tagged KLHL12 (**Figure 5D**), demonstrating that Kelch dimerization is sufficient to support BAY-805 induced USP21 recruitment. Interestingly, full length KLHL12 tagged with both HiBiT and LgBiT exhibited reduced interaction with USP21 in the presence of BAY-805 (**Figure 5D**), likely due to unfavorable tag orientation or conformational constraints imposed by BTB BACK-mediated dimerization. Together, these findings demonstrate that dimerization of Kelch domains is a critical determinant of BAY-805 induced USP21 recruitment to KLHL12.

Finally, we examined whether USP21 mutations we identified as disrupting BAY-805 binding (**Figure 4H)** also impaired USP21–KLHL12 association. Consistent with results from NalTSA and NanoBRET assays, mutations C398W, E311Y, E311F, and E366R abolished BAY-805 induced interaction between USP21 and KLHL12 (**Figure S5A**). Notably, E311F and E366R did not abolish USP21 catalytic activity (Figure S5B), suggesting that BAY-805-driven KLHL12 recruitment does not depend solely on USP21 inhibition.

### BAY-805 outcompetes ubiquitin from USP21 and enables KLHL12 binding

Superimposition of the BAY-805-bound USP21 model with the diubiquitin-bound USP21 crystal structure (PDB 2Y5B), ^31^ revealed close structural alignment (C_α_ RMSD = 0.351), implying minimal global conformational change upon ligand binding. However, a steric clash is evident between K48 of ubiquitin and the isobutyl group of BAY-805 (**Figure 6A-B**). Notably, the isobutyl group bearing analogue is the most potent compound in the BAY-805 series.^16^ Given the critical role of K48 in substrate ubiquitination and in positioning the K48-mediated isopeptide bond for cleavage by DUBs, our *in silico* model suggests that BAY-805 directly disrupts ubiquitin binding (**Figure 6B**).

**Figure 6.**
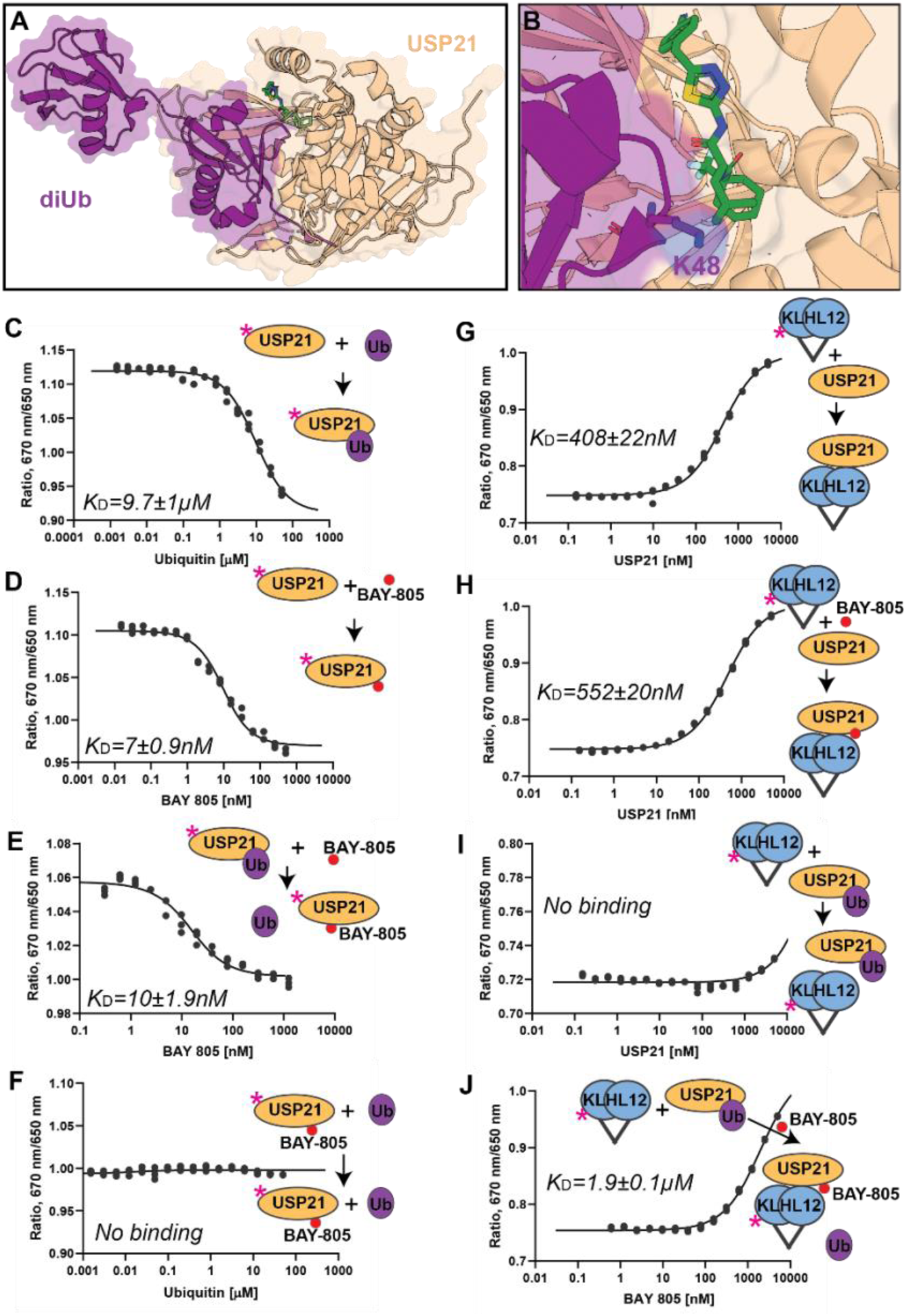
BAY-805 outcompetes ubiquitin and promotes USP21-KLHL12 interaction *in vitro*. (A) Structural overlay of the USP21-BAY-805 AlphaFold3 co-folding model with the USP21-diubiquitin experimental structure (PDB ID: 2Y5B). (B) Close-up view of (A) highlighting a steric clash between ubiquitin K48 and isobutyl group of BAY-805. (C) Binding of USP21 to ubiquitin measured by SpS, using USP21 labeled with Lys-reactive dye (magenta star) with titrated with increasing concentrations of ubiquitin. (D) Binding of USP21 to BAY-805, measured as in (C), with BAY-805 used as the titrant. (E) BAY-805 binds the USP21-ubiquitin complex with nanomolar affinity, as determined by titration of BAY-805 into labeled USP21 in the presence of excess ubiquitin. (F) Pre-incubation of labelled USP21 with 2.5 µM BAY-805 abolishes ubiquitin binding across a titration range of 0.0015 – 50 µM ubiquitin. (G-H) Interaction of USP21 with the C-terminally His-labeled recombinant HiBit-LgBit KLHL12 KELCH domain in the absence (G) or presence (H) of BAY-805, measured as a function of increasing USP21 concentration. (I) Ubiquitin (25 µM) disrupts USP21-KLHL12 interaction as assessed by titration of USP21 into the labeled KLHL12 KELCH domain in the presence of ubiquitin. (J) BAY-805 (2 – 5000 nM) restores USP21-KLHL12 interaction by displacing ubiquitin (15 µM Ubiquitin and 4 µM USP21). The C-terminally His-labeled labeled recombinant HiBit-LgBit KLHL12 KELCH domain was incubated with USP21 and ubiquitin, as in (I) and BAY-805 titrated All experiments in B-J were performed in triplicate.

We next experimentally validated this model using *in vitro* Spectral Shift (SpS) assays with recombinant USP21 protein and a His-tagged KLHL12 Kelch domain N-terminally fused to the HiBit and LgBit dimer complex, as full-length KLHL12 could not be purified in sufficient quantities (**Figure S6A-B**). The SpS binding assay measures changes in fluorescence emission of a protein labeled with an environmentally sensitive dye upon ligand binding.^32^ Titration of ubiquitin (0.0015 – 50 µM) into dye-labeled USP21 catalytic domain yielded a binding affinity of K_d_ 9.7 ± 1 µM, consistent with previous reports ^31^(**Figure 6C and S6C**). In contrast, BAY-805 bound USP21 with nanomolar affinity (**Figure 6D**), and this interaction was minimally affected by excess ubiquitin (25μM) relative to USP21 (10nM) (**Figure 6E**). However, preincubation of USP21 with BAY-805 effectively abolished subsequent ubiquitin binding (**Figure 6F**).

To assess USP21-KLHL12 interaction, we labeled the dimeric HiBit/LgBit-fused Kelch domains of KLHL12 (**Figure S6A-B**) and titrated USP21 (0.15 – 5000 nM). Surprisingly, the dimeric Kelch domains of KLHL12 bound USP21 in the presence and absence of BAY-805 with similar potency (**Figure 6G, H**). Kelch itself did not bind BAY-805 (**Figure S6D-E**). In contrast, addition of ubiquitin disrupted the USP21-KLHL12 interaction (**Figure 6I and S6F**). Notably, this disruption was reversed upon addition of BAY-805, which restored USP21-KLHL12 binding (**Figure 6K and S6G**). The competitive nature of the compound was supported in cellular co-transfections of ubiquitin variant Ubv21 with high affinity to USP21 ^33^ where Ubv21 blocked BAY-805 induced KLH12-USP21 association (**Figure S6H).** Collectively, these results support a model in which BAY-805 competes with ubiquitin at USP21 to enable KLHL12 binding (**Figure 6A-B**).

### BAY-805 induces USP21 colocalization with COPII defined by KLHL12 and SEC31

CUL3-KLHL12 facilitates the enlargement of COPII vesicles, a process required for secretory transport of large cargoes from the endoplasmic reticulum (ER) into the Golgi.^34–36^ To examine whether BAY-805 alters the subcellular localization of USP21 in relation to KLHL12 defined COPII structures, we performed immunofluorescence analysis in HeLa cells transiently overexpressing USP21 HaloTag (USP21 HT) and KLHL12 FLAG. Under control (mock treated) conditions, USP21 exhibited a largely diffuse cytoplasmic distribution, whereas KLHL12 displayed a punctate localization pattern, consistent with previous reports ^35^ (**Figure 7A**). Immunostaining of the endogenous COPII component SEC31A revealed numerous cytoplasmic puncta, a subset of which colocalized with KLHL12, consistent with enlarged COPII vesicles described previously.^20, 29, 35^

**FIGURE 7.**
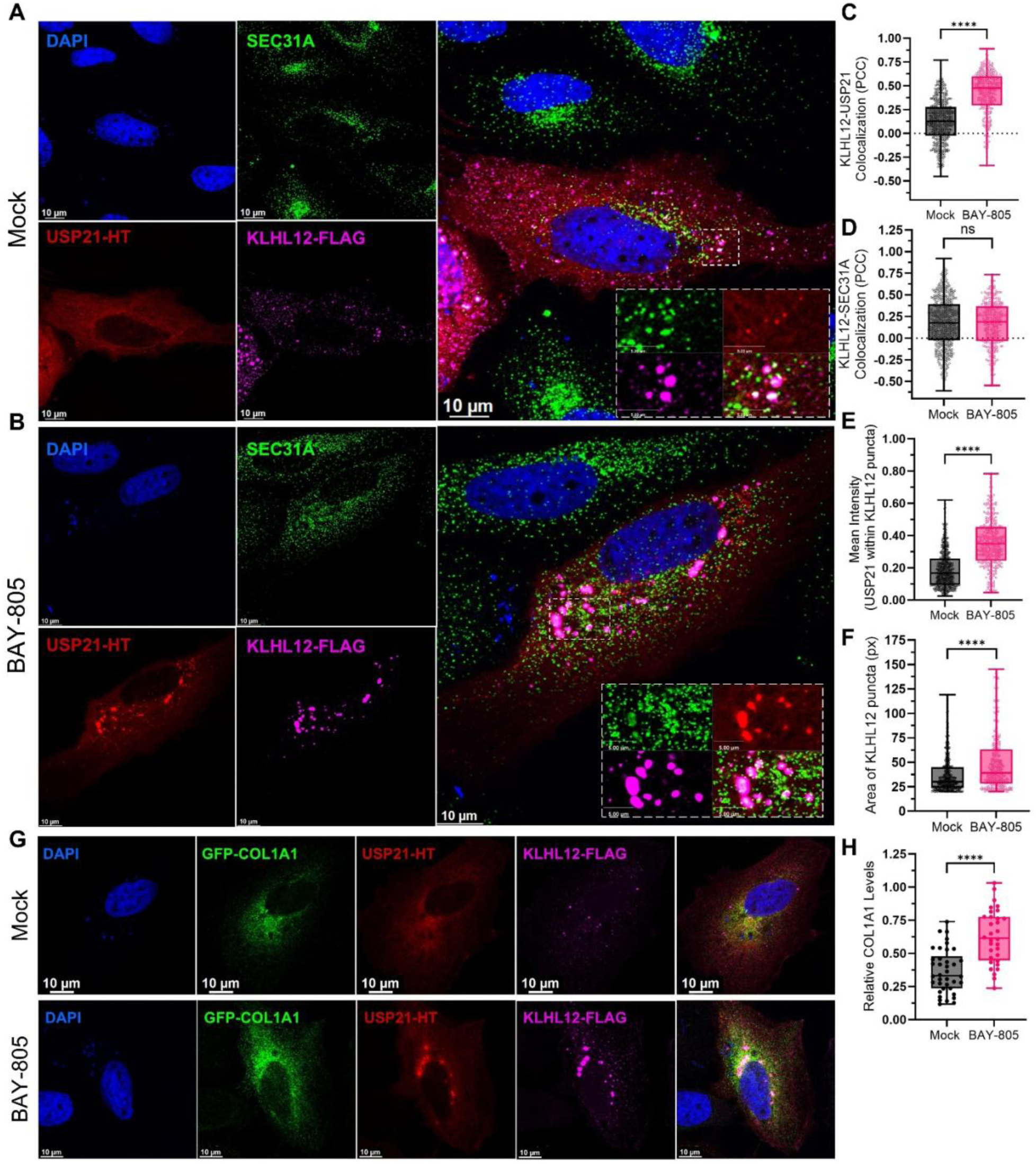
BAY-805 enables the USP21-KLHL12 association and recruitment of USP21 to COPII vesicles. (A-B) The localization of USP21-HT and KLHL12-FLAG in HeLa cells comparing to the COPII outer membrane subunit SEC31A in the (A) absence or (B) presence of BAY-805. (C-F) Cell Profiler analysis was conducted (*n* = 6-10 cells for each condition) across three biological replicates, of which a single replicate is shown. Colocalization by Pearson’s correlation coefficient (PCC) scores for: (C) KLHL12-USP21 and (D) KLHL12-SEC31A. (E) Signal intensity of USP21 within KLHL12 puncta, and (F) overall area of KLHL12 size were determined. (G) Trafficking of transiently overexpressed procollagen I (α1) (GFP-COL1A1) in HeLa cells with co-transfected KLHL12-FLAG and USP21-HT was determined in mock (*n* = 38 cells) and BAY-805 (*n* = 32 cells) treatment conditions. Representative images acquired on a Leica Stellaris 5 laser scanning confocal microscope from each experiment are shown. BAY-805 treatments were conducted for 30 minutes at 1 µM prior to cell fixing. (H) Quantification of relative intracellular COL1A1 levels. Asterisks (****) denote statistically significant findings for p<0.001; ns = not significant, combined across three biological replicates

Treatment with BAY-805 induced a marked redistribution of USP21 from a diffuse cytoplasmic pattern to punctate structures that colocalized with both KLHL12 and SEC31A (**Figure 7B**). Quantitative analysis revealed a statistically significant 3–4-fold increase in USP21– KLHL12 colocalization, as measured by Pearson’s correlation coefficient (PCC), upon BAY-805 treatment (**Figure 7C**). In contrast, BAY-805 did not significantly alter KLHL12–SEC31A colocalization (**Figure 7D**). Consistent with USP21 recruitment to KLHL12 defined COPII, BAY-805 increased the average area of KLHL12 positive COPII vesicles by approximately 30% and nearly doubled the USP21 signal intensity within these structures (**Figure 7E–F**).

To exclude the possibility that the HaloTag fusion affected USP21 localization, we confirmed that USP21 with a Myc tag displayed identical localization patterns under control and BAY-805 treated conditions (**Figure S7A**). Furthermore, the catalytically inactive USP21 mutant (C221A) that has been reported to increase the DUB enzyme affinity for ubiquitin^37^ failed to colocalize with KLHL12 upon BAY-805 treatment (**Figure S7B**). Consistently, NanoBRET assays showed that the catalytically inactive USP21 C221 mutant did not interact with KLHL12 in the presence of BAY-805 (**Figure S7C–D**). This mutant also failed to bind BAY-805 in cellular target engagement assays (**Figure S7E**). Interestingly, the E366R mutant of USP21, which did not bind BAY-805, was also not recruited to COPII by compound treatment (**Figure S7F–G).** Together, these data demonstrate that BAY-805 binding to USP21 is required to drive its recruitment to KLHL12-associated COPII vesicles

### BAY-805 disrupts the COPII trafficking of procollagen I

We next examined whether BAY-805 induced recruitment of USP21 to KLHL12 affects COPII-mediated transport of large secretory cargo. Procollagen trafficking from the ER is a well-established outcome of COPII vesicle enlargement and function,^38^ and CUL3–KLHL12 has previously been implicated in collagen export through this pathway.^20, 29^ To monitor collagen trafficking, we transiently overexpressed GFP-tagged procollagen I (COL1A1) and quantified total intracellular levels of unprocessed GFP COL1A1, thereby measuring ER accumulation prior to secretion. BAY-805 treatment resulted in an approximately 1.5–2.0-fold increase in intracellular COL1A1 levels (**Figure 7H**), indicating impaired procollagen export. To confirm the subcellular localization of GFP COL1A1, we performed co-staining with markers of the secretory pathway. GFP COL1A1 showed robust colocalization with the COPII marker SEC31A and localized proximal to the cis Golgi network, as indicated by GM130 staining (**Figure S8A–C**). GFP COL1A1 puncta defined by colocalization with SEC31A and/or GM130 are consistent with the identity of secretory cargo transitioning to Golgi ^39^ (**Figure S8D, arrows**). These findings are consistent with a model in which BAY-805 mediated recruitment of catalytically inhibited USP21 to KLHL12 contributes to disrupted COPII-dependent trafficking of large cargos such as procollagen

## DISCUSSION

Chemical inducers of proximity can rewire protein-protein interactions and reshape cellular signaling circuits.^40, 41^ While investigating USP21 interactors, we discovered that BAY-805 induced a robust association between USP21 and Kelch-like protein KLHL12, a substrate receptor of the Cullin3 ligase complex. Using a combination of LiP MS, in silico docking, and ΔHDX MS, we defined the BAY-805 binding site within USP21, showing that the compound engages the orthosteric S1 substrate binding pocket formed between the thumb and fingers domains. BAY-805 binding competes with ubiquitin and, in doing so, renders USP21 competent to interact with dimeric KLHL12. Strikingly, this chemically induced interaction was accompanied by pronounced colocalization of USP21 with KLHL12 positive COPII vesicles in cells. Together, these findings identify a novel proximity mechanism in which competition with the native ubiquitin substrate enables a new USP21 protein–protein interaction.

Previous studies have established that USP21 localizes to centrosomes and microtubule-associated complexes and regulates diverse processes including chromatin modification, transcription, centrosome function, and antiviral signaling. ^3, 5, 12, 13^ Our BioID analysis further revealed extensive USP21 proximity to microtubules, centrosomal components, and the homologous to augmin subunits (HAUS) complex, a key regulator of non-centrosomal microtubule organization.^42^ Notably, BAY-805 perturbed only a limited subset of USP21 interactions, with the most prominent effect being induction of the USP21–KLHL12 interaction, underscoring the specificity of this chemical perturbation.

KLHL12 functions as a CUL3 adaptor and promotes monoubiquitination of COPII coat components such as SEC31 and the co-adaptor PEF1, driving the enlargement of COPII vesicles required for transport of large cargoes, including collagen, from the endoplasmic reticulum (ER). ^20, 29, 35^ Substrate recognition by KLHL12 typically involves a conserved proline-rich “PGXPP” degron motif present in client proteins such as SEC31, PEF1, Disheveled proteins, PLEKH4, and Lunapark. ^23–25^ Although USP21 contains a unique insertion between conserved USP boxes 2 and 3 that harbors a proline-rich “PXXP” like motif, mutational analysis demonstrated that this motif is dispensable for both BAY-805 binding and KLHL12 interaction.

The association of USP21 with KLHL12, induced by BAY-805, prompted us to investigate how the recruitment of catalytically inhibited USP21 affects KLHL12-dependent COPII assembly and secretory cargo trafficking. COPII vesicles mediate ER to Golgi anterograde transport through assembly of a proteinaceous coat composed of SEC13/31 heterodimers. ^36, 43^ CUL3–KLHL12 facilitates vesicle enlargement required for transport of large cargoes such as collagen.^34–36^ We found that BAY-805 mediated recruitment of USP21 to KLHL12 increased the apparent size of KLHL12 positive COPII structures and impaired secretion of procollagen I. Notably, other DUBs, including USP8 and USP9X, regulate COPII size by deubiquitinating SEC31A and PEF1, respectively, and loss or catalytic inhibition similarly results in vesicle enlargement. ^20, 44^ Because BAY-805 mediated recruitment relies on ubiquitin competition, rendering USP21 inactive, and given that other DUBs (USP8, USP9X) regulate COPII vesicle size through deubiquitination of SEC31A and PEF1, the observed vesicle enlargement and procollagen accumulation may reflect contributions from both catalytic inhibition of USP21 on COPII substrates and the induced proximity to KLHL12. However, our findings on E311F and E366R mutations of USP21 that do not ablate catalytic activity argue that BAY-805 has distinct binding elements within the ubiquitin pocket that prevent ubiquitin binding, thus compromising catalytic activity and enabling other protein-protein interactions such as KLHL12. Further studies will be required to delineate the precise structural basis of USP21–KLHL12 engagement, providing insights into the BAY-805 catalytic inhibition vs USP21-KLHL12 association and how this interaction integrates USP21 function across cytoskeletal and secretory pathways.

Structural insights from docking and ΔHDX MS revealed that BAY-805 binds the S1 pocket of USP21 in a manner analogous to type III D USP7 inhibitors such as GNE 6776 and GNE 6640, which occupy the cleft between the α5 and α6 helices and sterically occlude ubiquitin binding. ^14, 45^ Despite substantial sequence divergence between USP21 and USP7, particularly in the insertion between conserved boxes 2 and 3 (residues ∼320–360), BAY-805 exploits a conserved architectural feature of USP catalytic domains. In USP21, this insertion comprises α5a helix (350-364), an acidic residue-rich turn, followed by α6a helix, absent in USP7 (nomenclature as in ^31^). Overall, surprisingly, despite structural differences between USP21 and USP7, the BAY-805 binding cleft in USP21 was similar to the USP7 binding GNE-6776 and GNE-6640, where ligands were positioned between the α5 and α6 helices of USP7, engaging D305. ^45^ In BAY-805, thiazole amine is predicted to establish a hydrogen bond with the carboxylate group of E366, while the aromatic ring participates in weak cation–π interactions with R365, and the thiazole amine establishes a hydrogen bond with the neighboring E366. This model is consistent with the ΔHDX-MS-observed decrease in deuterium uptake in the peptide spanning residues 363-373. The ΔHDX-MS also identified decreased deuterium uptake in peptide 389-397, which is a highly conserved β2 sheet that forms the ubiquitin-recognition “fingers”. It is possible that the Y362 and Q377 (β1 sheet) hydrogen bonding with BAY-805 that orients the ligand deep into a hydrophobic pocket, also distorts the β2 sheet position. However, this possibility needs further confirmation. It is important to note that our conventional HDX-MS approach, at a timescale of 0.25 to 10 mins, monitors exchange at the amide backbone, not residue sidechains. For these reasons, residues which form H-bonds through their sidechains in the AF3 docked model while leaving the backbone amide accessibility unchanged will not be captured. Similarly, there were also no statistically significant differences observed for the flexible loop (res. 323-338) at the timepoints tested, despite it being identified by LiP MS.

BAY-805 displays competitive behavior with ubiquitin, resulting in the USP21 form that interacts with KLHL12. Our SpS experiments support this, showing that in vitro BAY-805 does not appreciably alter the affinity between USP21 and KLHL12 but instead competes with ubiquitin. These findings are also consistent with cellular nanoBRET, where we observed a baseline USP21 and KLHL12 interaction that increased 4-5-fold with BAY-805 treatment. Interestingly, USP7 binding GNE-6776 or GNE-6640 was shown to sterically inhibit ubiquitin binding, locking USP7 in an inactive conformation. ^45^ Thus, BAY-805 acts by competing with the ubiquitin substrate and possibly increasing the cellular pool of USP21 that binds to KLHL12. More work is needed to decipher the structural mechanism of USP21-KLHL12 binding, and it should be noted that the existing empirical structures of USP21 are of ubiquitin- or ubiquitin variant-bound forms of USP21. ^6, 31, 33^ Thus, it is possible that the apo form of USP21 adopts a different folding state, as supported by structural shifts observed in HDX upon BAY-805 treatment. Overall, convergent evidence from HDX-MS, mutagenesis, and SpS competition supports BAY-805 binding at the orthosteric site and ubiquitin competition that underlies BAY-805-induced USP21-KLHL12 proximity.

## CONCLUSION

We identify a previously unrecognized chemical-induced proximity mechanism in which the USP21 inhibitor BAY-805 competes with ubiquitin, thereby enabling USP21 binding to the CUL3 adaptor protein KLHL12. In cells, BAY-805 bound USP21 is recruited to KLHL12 defined COPII vesicles, resulting in increased apparent size of KLHL12-positive COPII structures and impaired collagen trafficking. These findings reveal unanticipated novel USP21 interactions and highlight substrate competition as a powerful mechanism for chemically rewiring protein complexes. BAY-805 thus represents a valuable chemical probe to interrogate the dynamic functions of USP21 in ubiquitin signaling, vesicle trafficking, and cytoskeletal regulation.

## MATERIALS & METHODS

| REAGENT or RESOURCE | SOURCE | IDENTIFIER |
| --- | --- | --- |
| Antibodies |  |  |
| FLAG M2 | Sigma | F1804 |
| USP21 | Proteintech | 17856-1-AP |
| KLHL12 | Santa Cruz | sc-514874 |
| beta-Actin | Santa Cruz | sc-47778 |
| Anti-Myc tag rabbit | CST | 2278S |
| Anti-Myc tag mouse | CST | 2276S |
| Anti-LAMP1 rabbit | CST | 9091 |
| Anti-GM130 rabbit | CST | 12480 |
| Anti-FLAG rabbit | CST | 14793S |
| Anti-FLAG M2 mouse | Sigma-Aldrich | F1804 |
| IRDye 800CW (anti-rabbit, green) (1:4000) | LICORbio | 926-32213 |
| IRDye 800CW (anti-mouse, green) (1:4000) | LICORbio | 926-32212 |
| IRDye 680RD (anti-rabbit, red) (1:4000) | LICORbio | 926-68073 |
| IRDye 680RD (anti-mouse, red) (1:4000) | LICORbio | 926-68072 |
| Streptavidin IR800 (1:10,000) | LICORbio | 926-32230 |
| AlexaFluor 488 (anti-rabbit, anti-mouse) (1:500) | CST | 4408S (m), 4412S (r) |
| AlexaFluor 555 (anti-rabbit, anti-mouse) (1:500) | CST | 4409S (m), 4413S (r) |
| AlexaFluor 647 (anti-rabbit, anti-mouse) (1:500) | CST | 4410S (m), 4414S (r) |
| Bacterial and virus strains |  |  |
| Chemicals, peptides, and recombinant proteins |  |  |
| USP21 recombinant proteins | Generated in house | Methods below |
| Acetonitrile, Optima LC/MS Grade | Fisher chemical | A955-212 |
| Ammonium bicarbonate | Fluka | 09830-500G |
| Benzonase Nuclease | Sigma | E1014-25KU |
| DSS (disuccinimidyl suberate) | Thermo Scientific | 21555 |
| EDTA Disodium Salt 2-hydrate | AppliChem | A2937 |
| Formic acid | Sigma-Aldrich | 27001-500ML-R |
| HEPES | Sigma | H4034-500G |
| Iodoacetamide | Sigma | I1149-5G |
| Lysyl Endopeptidase, Mass Spectrometry Grade (Lys-C) | Wako | 121-05063 |
| iRT peptides | Biognosys | Ki-3002-2 |
| Phenylmethanesulfonyl fluoride (PMSF) | Sigma | P7626-5G |
| Protease Inhibitor Cocktail (PI) | Sigma | P8849 |
| Sodium chloride | Merck | 1.06404.5000 |
| Sodium fluoride | Sigma | S7920-500G |
| Sodium hydroxide pellets | Merck | 106482 |
| Strep-Tactin Sepharose resin, 50% suspension | iba | 2-1201-010 |
| Pierce TCEP-HCl | Thermo Scientific | 20490 |
| Tris - Hydrochlorid | AppliChem | A3452 |
| Triton X-100 | Sigma-Aldrich | T9284-100ml |
| Sequencing Grade Modified Trypsin | Promega | V511C |
| UltraMicroSpin Column | The Nest Group, Inc. | 800-347-6378 |
| Water HPLC Grade | Fisher Scientific | W/0106/17 |
| Sodium dodecyl sulfate | Sigma | 71729 |
| Urea | Fluka | 51459 |
| Vivacon 500, 30,000 MWCO Hydrosart, 100pc | Satorius | VN01H22 |
| 2-phospho-L-absorbic acid | Sigma-Aldrich | 49752-10G |
| (+)-Sodium L-ascorbate | Sigma-Aldrich | A7631-100G |
| Fluoroshield with DAPI | Sigma-Aldrich | F6057 |
| Paraformaldehyde 16% Aqueous Solution EM Grade | Electron Microscopy Sciences | 15710 |
| Critical commercial assays |  |  |
| NanoBRET | Promega |  |
| Deposited data |  |  |
| PRIDE proteomics data for BioID, LiP MS |  |  |
| The LiP MS proteomics data have been deposited to the ProteomeXchange Consortium via the PRIDE partner repository with the dataset identifier PXD080491 | PRIDE | PXD080491 |
| The BioID proximity MS data available via PRIDE | PRIDE | PXD041068 |
| HDX data | Zenodo | 10.5281/zenodo.22081973 |
| Experimental models: Cell lines |  |  |
| HEK293T<sup>REX</sup> | Invitrogen | R78007 |
| Hela | ATTC | CCL-2,<br>RRID:CVCL_0030 |
| U2OS | ATTC | HTB-96<br>RRID:CVCL_0042 |
| HEK293 | ATTC | CRL-1573<br>RRID: CVCL_0045 |
| Experimental models: Organisms/strains |  |  |
| Recombinant DNA |  |  |
| KLHL12-FLAG | This paper | N/A |
| USP21-HT | This paper | N/A |
| HT-KLHL12, KLHL12-HT | This paper | N/A |
| HT-CUL3 | This paper | N/A |
| HT alone | Promega |  |
| USP21-NL (WT, Δ323-366, Δ323-338, Δ331-345, Δ339-348, E366R, C398W, E311F, E311Y) | This paper | N/A |
| NL-KLHL12 (WT, Δ264-568, Δ132-568, Δ2-274, Δ2-131, Δ132-263) | This paper | N/A |
| HB/LB-KLHL12/Kelch, | This paper | N/A |
| HT-CTNNB1 | This paper | N/A |
| NL-DAPK1 | This paper | N/A |
| NL-KCTD6 | This paper | N/A |
| ANGEL1-HT | This paper | N/A |
| USP2-NL | This paper | N/A |
| USP22-NL | This paper | N/A |
| USP10-NL | This paper | N/A |
| NL alone | Promega |  |
| eGFP-hCOL1A1 | Addgene | #140110 |
| USP21-Myc (WT, C221A, C221S) | This paper | n/a |
| USP21-HT | This paper | n/a |
| KLHL12-FLAG (WT, and mutants) | This paper | n/a |
| USP21-BirA (N and Cterm) pcDNA5 FRT | This paper | n/a |
| Software and algorithms |  |  |
| CellProfiler v4.2.6 | Broad Institute | n/a |
| LAS-X Application Suite | Leica Microsystems | n/a |
| GraphPad PRISM 10 | GraphPad | n/a |
| Licor Odyssey F | LICORbio | n/a |
| Image Studio | LICORbio | n/a |

### BioID MS

#### BioID

Strep-Tactin Sepharose resin was crosslinked by incubation with 5 mM DSS in HNN buffer (50 mM HEPES (pH 8), 0.15 M NaCl, 50 mM NaF) at 37°C for 30 min. The reaction was quenched with 50 mM Tris-HCl at 37°C for 15 min. Both incubation steps were done at 37 °C and 1100 rpm. The beads were prepared a day ahead of the experiment.

The cell pellet of a 150 mm plate of T-Rex-HEK293 Flp-In cells (generated as below) stably expressing the miniTurbo-tagged USP21 was lysed with 1 mL RIPA buffer (50 mM Tris (pH 8), 0.15 M NaCl, 1% Triton, 1 mM EDTA, 0.1% SDS) supplemented freshly with 0.05% Benzonase, 1 mM PMSF, 0.2% PI. The lysate was briefly sonicated and incubated at 10°C for 20 min. The cleared lysate was incubated with 120 µL crosslinked beads (50% suspension) at 4°C on a rotary wheel for 1 h. After washing with 3x 1 mL RIPA buffer (without PMSF, PI and Benzonase), 3x 1 mL HNN buffer and 2x 1 mL ammonium bicarbonate (100 mM). The beads were transferred with 2x 200 µL ammonium bicarbonate (100 mM) into new tubes. After centrifugation at 200x g for 1 min the supernatant was removed, and the samples were denatured with 8 M Urea in ammonium bicarbonate (100 mM). After an incubation at 20°C for 10 min, the Urea was diluted to 4 M. The samples were reduced with 5 mM TCEP-HCl in ammonium bicarbonate (100 mM) for 30 min and 10 mM Iodacetamid for another 30 min. Both incubations were done at 37°C in the dark. The reduced proteins were digested with 0.5 ug Lys-C for 3 h. The samples were diluted to a final Urea concentration of 1 M with ammonium bicarbonate (100 mM) and digested with 1 ug Trypsin overnight. The peptides were separated from the beads by using 30 kDa Vivacon 500 spin column. The beads were washed with 200 µL and 100 µL 1 M Urea in ammonium bicarbonate (100 mM). The peptide solution was acidified with 5% formic acid and purified with equilibrated C18 UltraMicroSpin Column. The peptides were washed 3x with 5% acetonitrile containing 0.1% formic acid and then eluted with 2× 100µL 40% acetonitrile containing 0.1% formic acid. The eluted peptides were dried on a speed vacuum centrifuge and resuspended in 20 µL 2% acetonitrile containing 0.1% formic acid and 0.5x iRT peptides.

#### MS data dependent acquisition

LC-MS/MS analysis was performed on an Orbitrap Exploris 480 mass spectrometer (Thermo Scientific) coupled to an Easy-nLC 1000 system (Thermo Scientific). 1 µL of peptide solution was separated with a 120 min gradient from 3 to 30% acetonitrile at a flow rate of 300 nL/min on a column packed with 3 µm C18 resin. The data-dependent acquisition (DDA) mode was performed with the following parameters: one full FTMS scan (350 – 1’150m/z) at 120’000 resolutions followed by 20 MS/MS scans in the Ion Trap. Only charge states between or equal to 2 – 6 were included. An isolation window of 1.4 m/z was applied to isolate the selected ions. The normalized AGC target was set to 200% with a maximum injection time of 264 ms. The RF lens was set to 50%. The top 20 peaks were fragmented with a HCD of 30%. The MS2 spectra were measured with an Orbitrap resolution of 30’000 with isolation windows of 1.4 m/z. The AGC target was set to 200% with maximum injection time of 54 ms.

#### MS data independent acquisition

LC-MS/MS analysis was performed on an Orbitrap Exploris 480 mass spectrometer (Thermo Scientific) coupled to an Easy-nLC 1000 system (Thermo Scientific). 1 µL of peptide solution was separated with a 120 min gradient from 3 to 30% acetonitrile at a flow rate of 300 nL/min on a column packed with 3 µm C18 resin. The data-independent acquisition (DIA) mode was performed with the following parameters: one full FTMS scan (350 – 1’150m/z) at 120’000 resolutions. The normalized AGC target was set to 200% with a maximum injection time of 264 ms. The RF lens was set to 50%. The targeted MS2 spectra for the desired masses in the variable isolation windows listed below were acquired by fragmentation with a HCD of 30%. The Orbitrap resolution was 30’000 with variable scan ranges. The AGC was set to 200% with a maximum injection time of 66 ms and the RF lens was set to 50%.

#### MS Search Analysis

Hybrid libraries were generated using all relevant data independent acquisition runs and their respective data dependent acquisition runs of the pooled replicates, using the Pulsar search engine in Biognosys Spectronaut version 18. The standard settings were used, except that the digest rule was set additionally to LysC/P and the number of allowed missed cleavages to 4. The runs were searched in Biognosys Spectronaut version 18 against the generated library. The precursor-level data was exported and processed in protti version 0.6.0.9 (REF 36699412 PMID) in R version 4.3.1. The data was log2 transformed and normalized on the precursor level. Only precursors, which met the following condition were used for analysis: log2 intensity of 10 or higher, an absolute retention time variation of 0.35 min or lower to the predicted retention time and an elution group q-value of 1e^-3^ or lower for the interactome definition of USP21 or 1e^-4^ or lower for determination of the changes in proximity upon compound treatment. The protein abundance was calculated with the iq method, which considered only proteins with at least three peptides for quantification. The differential abundance was calculated, and significance was determined across the quadruplicate using a moderated t-test with the Benjamini-Hochberg multiple testing correction. For the identification of the changes in proximity upon compound treatment, we considered proteins with an absolute log_2_(fold change) of 1.0 or higher and an adjusted p-value of below 0.05 as significantly changed. For the identification of the interactome, we considered proteins with a log_2_(fold change) of higher than 2.0 compared to miniTurbo control and an adjusted p-value of below 0.01 as significantly changed. Only the two conditions, which were compared, were analyzed in R together. For identifying the interactome we further considered only proteins, which were seen in less than 20% of the experiments in the CRAPome database. We then did a GO terms enrichment for components with Gorilla (REF 19192299, 17381235) by using two unranked lists of genes. The target list contained all significantly changed proteins of the comparison of USP21 tagged with miniTurbo at the C- or N-terminus to miniTurbo only cells. The background list contained all proteins from those experiments which were quantified in at least one condition twice.

### Identification of differential structural protease susceptibility of USP21 in the presence/absence of BAY-805 with LiP MS

#### Sample Preparation

Purified USP21 was diluted in LiP buffer (100 mM HEPES pH 7.5, 150 mM KCl, 1 mM MgCl2) to 0.04 μg/ μl for a total protein amount of 2 μg per sample in a volume of 50 μl. BAY-805 or the negative control compound (10 μM, 1 μM, 100 nM, 10 nM, 1 nM, 100 pM, 10 pM) or vehicle was added to the USP21 samples in triplicates and incubated for 5 minutes at 25°C. Subsequently, proteinase K (PK) (0.02 μg) was added and incubated for 5 minutes at 25°C. The PK was then inactivated by incubation at 99°C for 5 minutes. Finally, the samples were placed on ice for 5 minutes. The lysate was transferred to a new Eppendorf tube and diluted 1:2 with 10% DOC (56 μl) for a final concentration of 5% DOC. The disulfide bonds were reduced with 5 mM TCEP and incubated for 40 minutes at 37°C, 800 rpm in a thermomixer. The thiols were alkylated with 40 mM IAA and incubated for 30 minutes at 30°C in the dark, at 800 rpm. The samples were diluted 1:5 with 100 mM ABC to a DOC concentration of 1%. Lys-C (0.5 μg) and trypsin (0.5 μg) were added, mixed and incubated overnight at 37°C and 800 rpm. The digestion was stopped by adding 50% formic acid (FA) to reach a final concentration of 2% and shaken at 800 rpm at RT for 10 minutes. Precipitated DOC was removed by filtration using 0.2 µm PVDF filter plates (Corning) on a vacuum manifold. C18 columns were washed with 100 µl MeOH and 2× 100 µl Buffer B (50% ACN, 0.1% FA), equilibrated with 3× 100 µl Buffer A (0.1% FA). The samples were transferred onto columns, washed with 3× 100 µl Buffer A and eluted 2 times with 50 µl Buffer B. The samples were dried under vacuum at 45°C. The samples were immediately resuspended in 20 µl Buffer A, shaken at 800 rpm for 5 minutes and sonicated for 5 minutes. The samples were centrifuged at 18000x g for 15 minutes. Of the supernatant, 9 μl were transferred to the MS vials. To the samples, 1 μl iRT peptide mix (Biognosys) prepared according to manufacturer protocol and diluted 1:2 in Buffer A was added. Library samples were generated by combining all replicates in a single sample.

#### Measurements

Samples were measured on an Orbitrap Fusion Lumos fitted with a Waters Acquity LC system. Peptides were separated on a 40 cm column packed with ReproSil-Pur C18-AQ 1.9 µm beads (Dr. Maisch) using a 120-minute gradient (3% to 35% ACN with 0.1% FA). Library samples were measured in DDA, and all samples were measured in DIA. For the DDA measurement, the MS1 spectra were acquired in a scan range of 350-1150m/z with an Orbitrap resolution of 120000. The normalized AGC target was set to 200% with a maximum injection time of 100 ms. The RF lens was set to 30%. The targeted MS2 spectra were acquired for the top 20 peaks in the spectrum and fragmented with a HCD of 30%. The spectra were measured with an Orbitrap resolution of 30000 with isolation windows of 1.6 m/z. The AGC target was set to 200% with a maximum injection time of 54 ms. For the DIA measurements, the MS1 spectra were acquired in a scan range of 350-1400 m/z with an Orbitrap resolution of 120000. The normalized AGC target was set to 50% with a maximum injection time of 100 ms. The RF lens was set to 30%. The targeted MS2 spectra were acquired for the desired masses with variable isolation windows listed below and fragmented with a HCD of 28%. The spectra were measured with an Orbitrap resolution of 30000 with variable scan ranges. The AGC target is set to 200% with a maximum injection time of 54 ms. The RF lens was set to 30%.

#### MS Data Evaluation

Library samples were searched using Spectromine version 3.2 with BGS factory settings, except that semi-specific peptides were included in the search. The verified protein entries of all human proteins on Uniprot were included in the search space. This library was used to search the DIA files in Spectronaut version 16.2. The BGS factory settings were modified to filter the data by q-value during the quantification. The data were exported and analyzed with protti 0.5.0. Peptides were filtered for a raw intensity > 10 and for proteotypicity. Dose response curves were fit and filtered for curves that have a correlation > 0.85. Additionally, anova testing across the intensities measured at each probe concentration for each peptide was performed and the p-values were adjusted using the method of Benjamini Hochberg. Only curves with adjusted p < 0.05 were considered significant.

### Vectors and cloning

All cloning was performed using in-fusion HD EcoDry Mix (Takaro 639689) using pcDNA5FRT (Invitrogen), pNF C and N NanoLuc, HaloTag HT (Promega) vectors. Mutagenesis was performed using Q5 site directed mutagenesis kit (NEB) according the manufacturers instructions.

### Tissue culture

HEK293, HEK293 T-Rex, HeLa and U2OS cells were grown in high glucose DMEM containing 1 mM sodium pyruvate, 1× antibiotic-antimycotic reagent and 10% fetal bovine serum. Cells were kept in a 5% CO2, 37 °C incubator and tested for mycoplasma presence.

### Generation of stable HEK293 Flp-In T-REx cells for BioID

For each experimental replicate, HEK293 Flp-In T-REx cells were co-transfected with pOG44 (Flp-recombinase expression vector) and plasmid (pcDNA5) encoding for 3xFLAG-USP21 N and C tagged contructs, or FLAG-miniTurbo. Transfections were performed with X-tremeGENE™ HP DNA Transfection Reagent according to the manufacturer’s instructions. Following transfection, cells were selected with hygromycin at 200 µg/ml.

### NanoBRET

The interaction nanoBRET was performed as described before.^46^ HEK293T cells were seeded in 96-well white plates (4 × 104 cells/well, Greiner) and co-transfected with 0.001 µg/well NL-constructs, 0.03 µg/well HT-constructs, and 0.07 µg/well empty vector using XtremeGene HP transfection reagent, following the manufacturer’s instructions. The next day the media was replaced with DMEM (no phenol red) supplemented with 4% FBS, 100 U/mL penicillin, and 100 μg/mL streptomycin, with or without 1000-fold diluted HaloTag® NanoBRET™ 618 Ligand (Promega) and incubated with BAY-805 or DMSO control for 1-4 h. Next, 100-fold diluted NanoBRET™ Nano-Glo® Substrate (Promega) was added and donor emission at 460 nm and acceptor emission at 618 nm were read within 10 min of substrate addition using a ClarioStar plate reader (Mandel Scientific). For kinetic NanoBRET measurement, the next day after transfection, media were replaced with CO2-independent media (Leibovitz’s L-15 no phenol red, Wisent), supplemented with 4% FBS, 100 U/mL penicillin,100 μg/mL streptomycin, +/- 1000-fold diluted HaloTag® NanoBRET™ 618 Ligand and 100-fold diluted Intracellular TE Nano-Glo® Vivazine® Substrate (Promega). After 2h of equilibration at 37 °C, the Bay-805 compound was added, the plate was sealed with Breathe-Easy Sealing Film (Diversified Biotech), and donor emission at 450 nm and acceptor emission at 618 nm were read immediately every 15 min for 4 h at 37 °C using a ClarioStar plate reader. The NanoBRET ratio was determined by subtracting 618/460 (acceptor/donor) signal from cells without NanoBRET™ 618 HaloTag Ligand × 1000 from 618/460 signal from cells with NanoBRET™ 618 Ligand × 1000. For UbV21 experiments, HA-Ub or UbV21 ^33^ were cotransfected.

### NalTSA

HEK293T cells were seeded in 6-well plates at 1 × 106 cells/well in DMEM supplemented with 10% FBS, 100 U/mL penicillin, and 100 μg/mL streptomycin. Cells were transfected with 0.2 μg/well of C-terminally NL-tagged USP21 constructs together with 1.8 μg/well empty vector using X-tremeGENE HP transfection reagent, according to the manufacturer’s instructions. The following day, cells were trypsinized, pelleted by centrifugation, and resuspended in Opti-MEM without phenol red at a density of 5 × 105 cells/mL. Cell suspensions were dispensed into white 384-well PCR plates (Axygen) at 20 μL/well, with one well allocated per temperature point (37– 60 °C). Plates were heated for 3 min in a Mastercycler X50h thermocycler, followed by cooling to room temperature (22 °C) for 2 min. Subsequently, 10 μL of Opti-MEM containing 167-fold diluted NanoBRET Nano-Glo Substrate and 500-fold diluted extracellular NL inhibitor (Promega) was added to each well. Bioluminescence signals were measured using a CLARIOstar plate reader. Data were normalized to the 37 °C data point and fitted using nonlinear regression analysis in GraphPad Prism 11 to determine apparent melting temperatures (Tm) values.

### Immunoprecipitation

Immunoprecipitation was performed to assess the increased interaction between USP21 and KLHL12 induced by BAY-805. In 6 cm dishes, 1.2 × 106 HEK293T cells was seeded, and reverse transfection was conducted with 2 µg KLHL12-FLAG and 2 µg USP21-HT constructs using X-tremeGENE™ HP DNA Transfection Reagent (Sigma) following manufacturer’s protocol. BAY-805 (1 µM) or DMSO was added to cells 48 h post-transfection for 2-4 h. Cells were then harvested and lysed with cell lysis buffer for 15 min on ice: 50mM Tris-HCl (pH 7.5), 150mM NaCl, 1% Triton X-100, 0.1% Na deoxycholate, supplemented with protease inhibitors (Aprotinin 0.25 µg/ml, Leupeptin 0.25 µg/ml, Pepstatin A 0.25 µg/ml, E-64 0.25 µg/ml), 1 mM N-Ethylmaleimide (NEM), 1 mM AEBSF, and benzonase. Lysed cells were then centrifuged at 13,000 rpm to collect supernatant, which were diluted two-fold for immunoprecipitation. Diluted cell lysates were added to Magne® HaloTag® Beads (Promega) together with BAY-805 or DMSO, and incubated overnight at 4 ⁰C. Gel-loading buffer was used to elute the beads, followed by western-blot analysis with indicated antibodies. RIPK1 ubiquitination experiments were performed as before ^33^ in HEK293 cells with 1.5 ug for HA-Ub and 2ug FLAG-RIPK1, 0.5 ug for USP21-WT, 0.3 ug USP21-C221S, 1 ug for USP21-E311F and 1 ug for USP21-E366R transfections, lysis and antiFlag precipitations as above.

### Detection of KLHL12 and USP21 localization in cells using immunofluorescence

Determination of the effect of BAY-805 treatment on KLHL12, USP21, and SEC31 subcellular localizations was conducted in HeLa cells using confocal microscopy. In brief, 8.0 × 10^4^ cells were plated on 18-mm cover glass (VWR, no.1) in a 12-well plate. The subsequent day, cells were co-transfected with overexpression plasmids for USP21 C-terminally fused with HaloTag (USP21-HT), and KLHL12 C-terminally fused with 1x FLAG (KLHL12-FLAG) using X-tremeGene HP (Sigma-Aldrich) according to the manufacturer’s instructions, and approximately 4-6 hours later the media was replaced. The next day, the cells were treated with HT 618 ligand (Promega) according to the manufacturer’s instructions for rapid labeling. Briefly, the media was replaced with media containing HT 618 ligand (1:1000) and incubated for 30 minutes. Subsequently, the cells were washed once once with fresh media and incubated for an additional 30 minutes to washout the unbound ligand. During this 30-minute washout period, BAY-805 (1 μM) or DMSO was added to the cells. The media was then removed, washed with PBS, and the cells were fixed with 4% PFA diluted in PBS for 15 minutes. Afterwards, the PFA was removed and the cells were permeabilized with 0.1% Triton-X-100 diluted in PBS for 15 minutes, and then blocked with 2% BSA diluted in PBS-T (0.1% Tween-20) for 1-hour. Between each of the steps after fixing, a single wash of PBS was performed. The cells were incubated with primary antibody diluted in 2% BSA in PBS-T overnight at 4°C with rabbit anti-FLAG (CST #14793, 1:1000), and mouse anti-SEC31A (BD Transduction Labs, 612351, 1:500). The next day, primary antibody was removed and washed with PBS (5 minutes x 3), incubated with secondary antibody at room temperature for 1-hour (mouse AlexaFluor 488 and rabbit AlexaFluor 647, 1:500). The cells were then washed with PBS (5 minutes x 3) and the cover glass was mounted to the slide with Fluoroshield medium containing DAPI (Sigma-Aldrich).

#### Image acquisition using confocal microscopy

Microscopy was performed on a Leica Stellaris 5 confocal laser scanning microscope. Images were acquired sequentially (between lines) with the following laser lines, 405, 488, 561, and 638, using HyD S detectors. The images were acquired at 512×512 resolution (400 hz) with a 63x oil objective at zoom factors 2-4x. Monitored channels: scan 1 - DAPI (ex. 405 nm, emi. 425-497 nm), scan 2 - SEC31A/GFP-COL1A1 (ex. 488 nm, emi. 497-556 nm), scan 3 - USP21-HT (ex. 561 nm, emi. 566-643 nm), and scan 4 - KLHL12-FLAG (ex. 638 nm, emi. 643-792 nm). CellProfiler (4.2.6) was used for all downstream analysis of images which included the identification of relevant puncta (objects) where stated (eg. KLHL12), and the corresponding quantitation of signal intensities, colocalization scores, and puncta sizes from the identified objects were determined. The CellProfiler pipelines were manually optimized to ensure that the objects were appropriately identified each experimental condition. Statistics and plots of the analyzed data were generated using GraphPad Prism 10.

##### Procollagen trafficking using immunofluorescence

To visualize the effect of BAY-805 on the trafficking of procollagen (type 1, chain α1), HeLa cells were cotransfected with USP21-HT, KLHL12-FLAG, and GFP-COL1A1. Approximately 4-6 hours later, the media was replaced and ascorbate (50 µg/mL) and 2-phospho-L-ascorbate (2 mM) were added to the media. The GFP-COL1A1 plasmid was kindly gifted by Sergey Leikin (Addgene #140110). COL1A1 bears the N-terminal N2 propeptide cleavage site which spans the GFP-COL1A1 fusion, and therefore proteolytic processing upon COL1A1 secretion leads to loss of the GFP signal in the extracellular matrix. To normalize expression levels of each individual construct, CellProfiler (4.2.6) was used to calculate total COL1A1 signal intensities for each cell and were taken as a ratio of the total levels of USP21 and KLHL12. Only cells of which contained low to moderate expression levels were chosen for imaging, taking care to avoid cells which contained large COL1A1 aggregates due to overexpression. Transfection times were conducted to not exceed 16-18 hrs to avoid excessive COL1A1 levels. Subsequent fixation, permeabilization, blocking, and antibody treatments were conducted as previously described.

##### Protein production

The human USP21 (Q9UK80, 209−563) with an N-terminally fused thrombin cleavable Hexa-His Tag was expressed in E. coli BL21(DE3) following 0.25 mM isopropyl β-D-1-thiogalactopyranoside (IPTG) induction at 17 °C overnight. For purification, bacterial supernatant was applied to Protino Ni-NTA beads, washed with buffer A, and eluted with buffer B (buffer A with 300 mM imidazole) using a linear gradient. The elution pool was diluted 1:10 with buffer C (20 mM Tris pH 7.5, 10% glycerin, 1 mM TCEP), filtered for application to a preequilibrated MonoS 10/100 GL (Cytiva) cation exchange column. The bound protein was eluted by running a linear gradient from the low salt buffer (buffer C + 50 mM NaCl) to the high salt buffer (buffer C + 1000 mM NaCl). The cation exchange chromatography step was applied and protein purified by size exclusion chromatography using a HiLoad 26/600 Superdex 200 pg column in buffer D (20 mM Tris pH 7.5, 150 mM NaCl, 10% glycerin, 1 mM TCEP). KLHL12 Kelch domain (274-568) with LgBit or HIBit chimeric constructs in pFBOH-MHL-His were expressed in Sf9 cells and purification performed similarly to above.

##### HDX-MS

Samples of USP21 catalytic domain res. 209-563 (7.5 uM) with and without BAY805 (7.5 uM or DMSO-only) were prepared in 20 mM TRIS (pH 7.5), 50 mM NaCl, 0.01% Triton X-100. With a 2:15 dilution into deuterium buffer (10 mM phosphate buffer, 150 mM NaCl, pD 7.5), samples were labeled for 0.25, 1, and 10 mins at 20 °C. Upon quenching the labeled sample 1:1 in 100 mM phosphate buffer, pH 2.5 for 2 min at 0 °C, the samples were injected onto a 1:1 Nep2-Pep (AffiPro) digestion column at 15 °C, followed by desalting (ACQUITY UPLC BEH C18 VanGuard Precolumn, Waters Corp.) and reverse phase separation (ACQUITY UPLC BEH C18 Column, Waters Corp.) at 0 °C. Eluting peptides were electrosprayed and analyzed by Select Series Cyclic IMS-MS (Waters Corp.) set to HDMSe mode. Peptides were identified using ProteinLynx Global Server 3.0.3 (Waters Corp.) and the HDX-MS was analyzed using DynamX 3.0 (Waters Corp.). Results were visualized using PDB 2Y5B (REF 21399617) in PyMOL^TM^ 3.1.4.1 (Schrodinger, LLC) and GraphPad Prism 10.3.0 (507). For sequence coverage and redundancy, see Supplementary figure 4. To be considered statistically significant, the cumulative HDX differences of a peptide over the three timepoints needed to exceed triple the cumulative propagated error (each timepoint was collected in technical triplicate). ^47^

##### Co-folding with AlphaFold-3, Boltz-2 and Chai-1

The complex structure of the USP21-BAY-805 was predicted from the amino acid sequence of the USP1 catalytic domain (residues 210-565) and the SMILES string of BAY-805 using **AlphaFold-3 (AF3)**, Boltz-2, and Chai-1. In the case of AlphaFold-3, the model was run using five seeds to generate a single best-scoring model based on the integrated confidence metrics (ranking score) while for Boltz-2 --use_msa_server and --usepotentials settings (to allow multiple sequence alignment and inference-time steering potentials, respectively) were enabled to sample five structures, using 10 recycling steps and the default of 200 reverse diffusion steps. Similar to Boltz-2, five sample structures were generated using the default settings of Chai-1.

##### In silico mutations

Mutations were first generated and evaluated using the built-for-purpose feature available in ICM (Molsoft, San Diego). This package independently computes the changes in protein stability and binding free energy of a protein-ligand complex upon mutation of a single residue while keeping the rest of the protein structure rigid. A positive energy value for protein stability indicates that the mutation is likely leading to destabilization. The stability of the system (with and without ligand) was then evaluated along 100 nanoseconds molecular dynamics simulations conducted with ICM and open-MM forcefield.

##### Spectral shift

Spectral shift requires labeling of a target protein with a fluorophore using a variety of coupling chemistries. Here, USP21 was labeled using a lysine-reactive NHS labeling technology where the fluorescent dye was covalently attached to a surface-exposed lysine residue by following the supplier-provided labeling protocol (Cat # NT-L021; NanoTemper Technologies GmbH). The final labeled USP21 concentration used in the assay mixture was 10 nM. The KLHL12 (both N and C-terminal His^6^-tagged versions) labeling was performed through site-specific affinity labeling using a His-tagged target labeling kit (Cat # NT-L128; NanoTemper Technologies GmbH). In short, the His-tagged proteins were diluted in the assay buffer (20 mM HEPES, pH 7.5, 150 mM NaCl, 0.005% Tween-20, and 1 mM DTT) and set up a 16-point serial dilution with a top concentration of 1µM of target protein in 10 µl assay buffer for each point in a 384-well plate. An equal volume of the 4 nM labeling dye dissolved in the assay buffer was added to the protein wells. The protein and the dye mix in a 20 µl reaction volume were incubated at room temperature in the dark for 10 minutes. The affinity between the His-tagged proteins and the labeling dye was measured using the spectral shift method. The final His-tagged labeled protein concentration in the assays was determined based on the observed *K*_D_ between the target protein and the labeling dye. For interaction analysis, the compounds/protein analytes were dissolved in the assay buffer at a varying analyte concentration (for dose-response titration) by keeping 4% DMSO (2% final after mixing with protein mix). The labeled protein mix was prepared by adding 100 nM protein (final 50 nM) and 4 nM (final 2 nM) dye in the assay buffer. The protein mix was then incubated for 10 min at room temperature for target labeling. 10 µL of each of the compound/protein analyte mix and the labeled protein mix were mixed into the assay wells in a 384-well assay plate. The assay mixture was then incubated for 30 minutes at room temperature before measuring the emission intensities using the Dianthus instrument equipped with dual-emission (650 nm and 670 nm) detection optics (NanoTemper Technologies GmbH). The affinity (*K*_D_) or the displacement (*K*_disp_) of the compound/protein analyte was calculated by plotting the ratios of the fluorescence intensities at two fixed wavelengths as a function of the compound/protein analyte concentration and fitting the data to a 1:1 binding model using DI.Screening Analysis v2.2 software.

## ACKNOWLEDGEMENT

We thank Dr Fabian Goricke, for BAY-805 binding modeling analysis. We thank Dr Alex Bullock and Dr Evmorfia Dalietou, University of Oxford for providing their expertise on KLHL12. M.K. Q.L. E.W. were funded by a MITACS Elevate fellowships. D.B.L and R.J.H. is supported by funding from CIHR, NSERC, CFI. D.B.L. was supported by Cancer Research Society and Leukemia and Lymphoma Society of Canada. R.J.H is supported by the Hereditary Disease Foundation, and the Connaught Fund. S.K, V.R. and M.G. were supported by EU/EFPIA/OICR/McGill/KTH/Diamond Innovative Medicines Initiative 2 Joint Undertaking [EUbOPEN grant 875510]. Structural Genomics Consortium is a registered charity (no: 1097737) that receives funds from Bayer AG, Boehringer Ingelheim, Bristol Myers Squibb, Genentech, Genome Canada through Ontario Genomics Institute [OGI-196], Canada Foundation for Innovation Ontario Research Fund, MITACS, EU/EFPIA/OICR/McGill/KTH/Diamond Innovative Medicines Initiative 2 Joint Undertaking [EUbOPEN grant 875510], Janssen, Merck KGaA (aka EMD in Canada and US), Pfizer and Takeda.

## SUPPLEMENTARY DATA

**Figure S1. (Relating to Fig 1).**
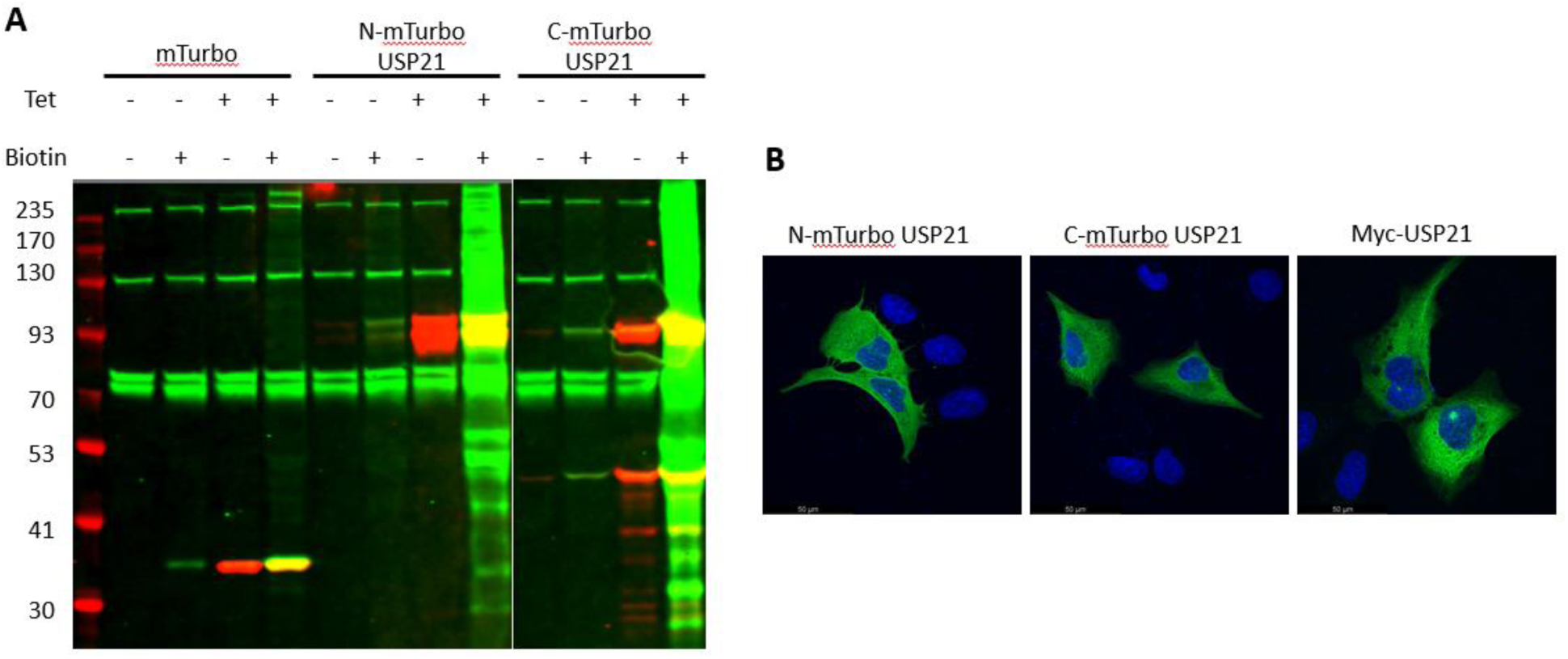
Characterization of USP21-mTurbo expression, inducibility, and localization. (A) Western blots for T-REx HEK293 cells and upon tetracycline induction and addition of biotin. Flag immunoblotting (red) and streptavidin detection of biotinylated proteins (green). (B) Immunofluorescence in transiently transfected U2OS cells demonstrating similar localization of mTurbo tagged USP21 as compared with Myc tagged USP21, detection anti-Flag or anti-Myc (green), DAPI nuclei staining (blue), panel scale is 50μm.

**Figure S2. (Relating to Fig 2).**
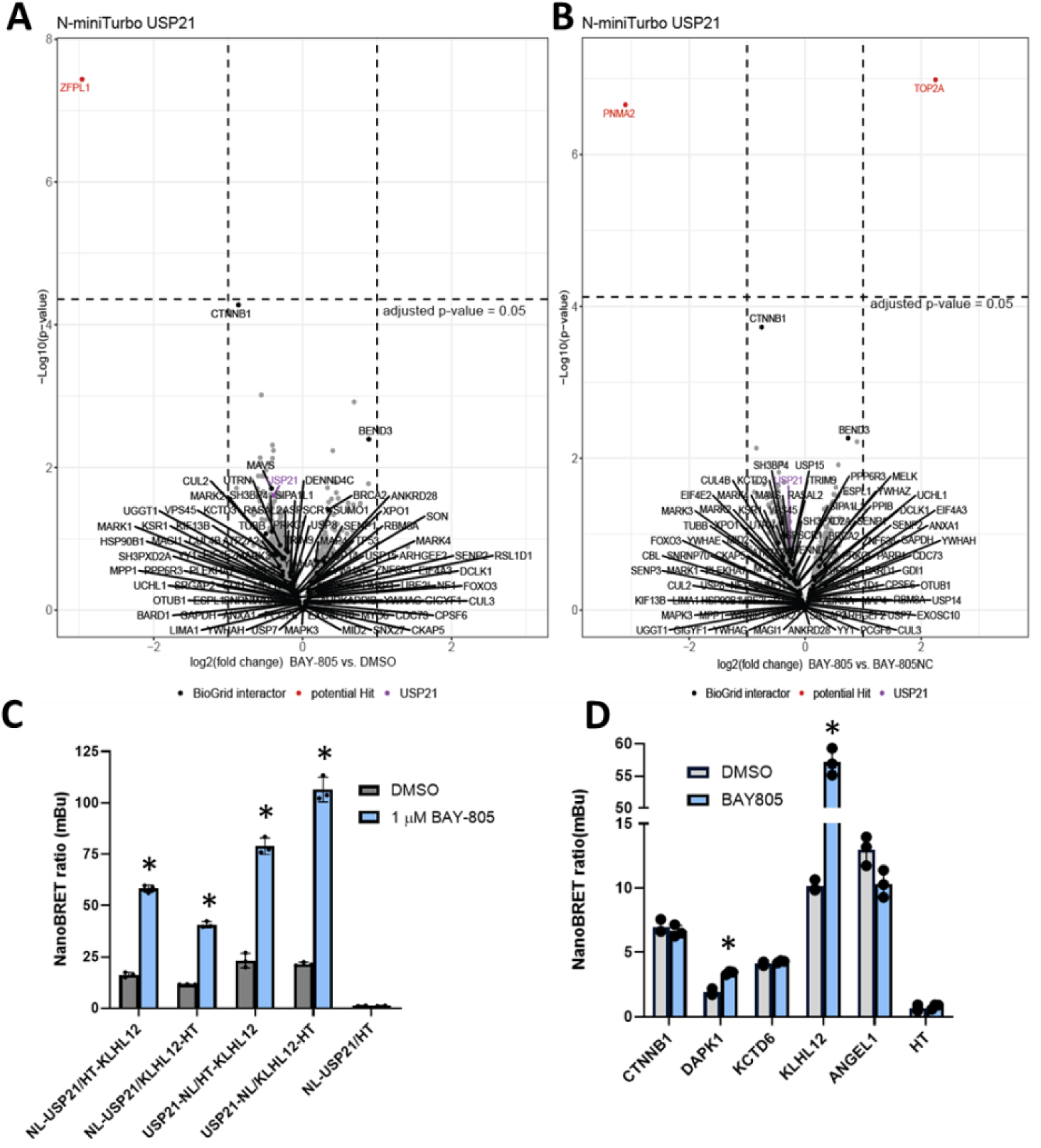
USP21 and KLHL12 association elicited by BAY-805. (A-B) BAY-805 effects on N-terminally tagged USP21 BioID. USP21 proximity profiling of HEK293 cells expressing N-tagged miniTurbo-USP21. Volcano plot displaying the log2 fold changes (x-axis) and the log10 p-values (y-axis) for proteins of the USP21 proximity in the presence of BAY-805 (1µM) compared to DMSO. Significance cut offs are indicated by dashed lines (log2FC ≥ |1| and adjusted p-value ≤ 0.05). (C) Validation of USP21-KLHL12 association by nanoBRET by using different tag orientations. BAY-805 induced significant increase is denoted by *, p<0.05. (D) Validation of other BAY-805 driven interactions. BAY-805 induced significant increase is denoted by *, p<0.05.

**Figure S3 (Relating to Fig 3).**
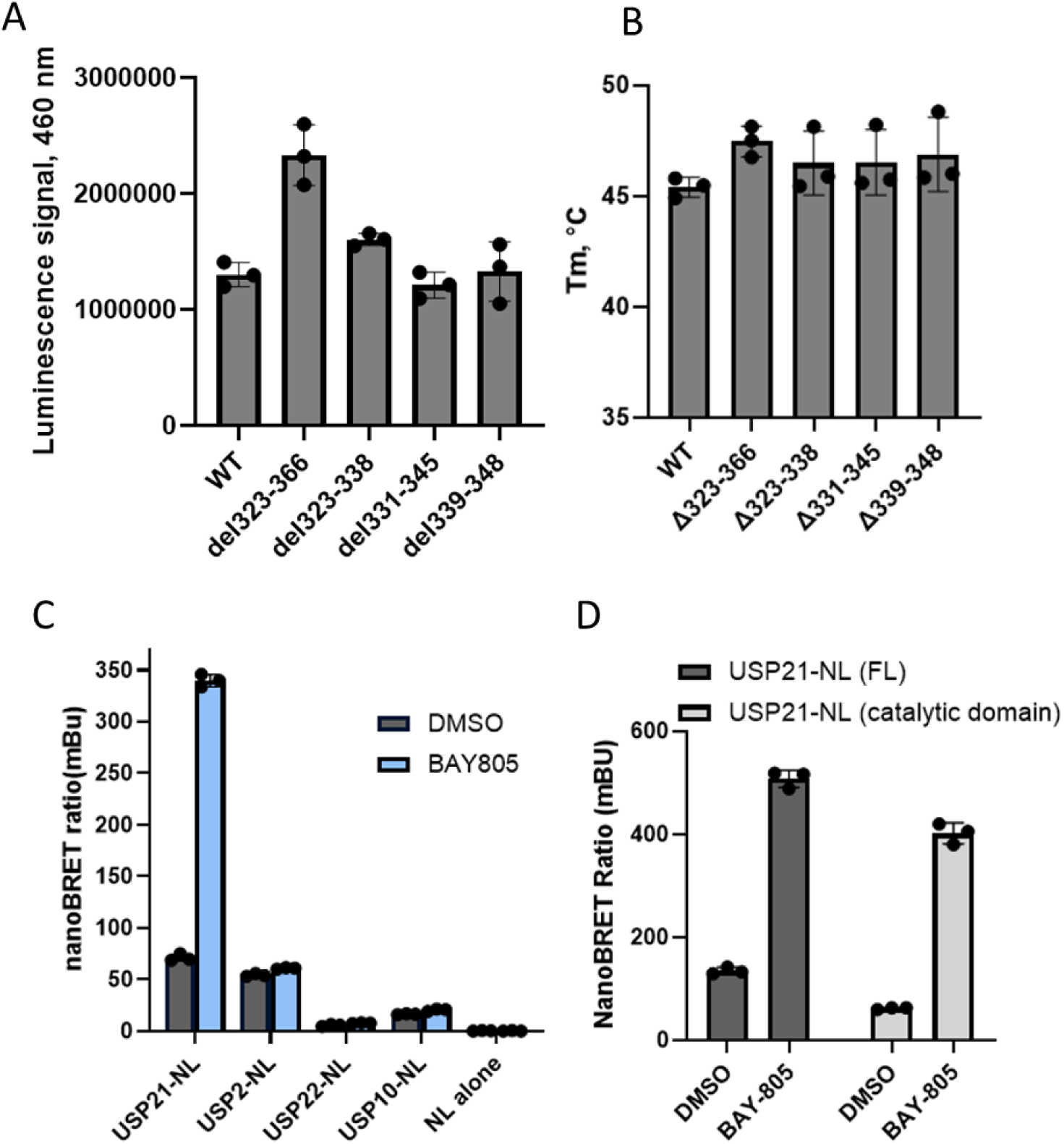
USP21 regions driving BAY-805 binding and association with KLHL12. (A) The levels of USP21 truncations used in Fig 3C-D to interrogate BAY-805 binding. The levels are inferred from USP21-nLuc luminescence. (B) Thermostability of USP21 mutants used in Fig 3C-D (C) BAY-805 increases KLHL12 association with USP21 but not USP2, USP22 or USP10. NanoBRET was performed as in Fig 2D. (D) Catalytic domain of USP21 is sufficient for BAY-805 (1 μM) induced KLHL12 association. NanoBRET was performed as in Fig 2D.

**Figure S4 (Relating to Fig 4).**
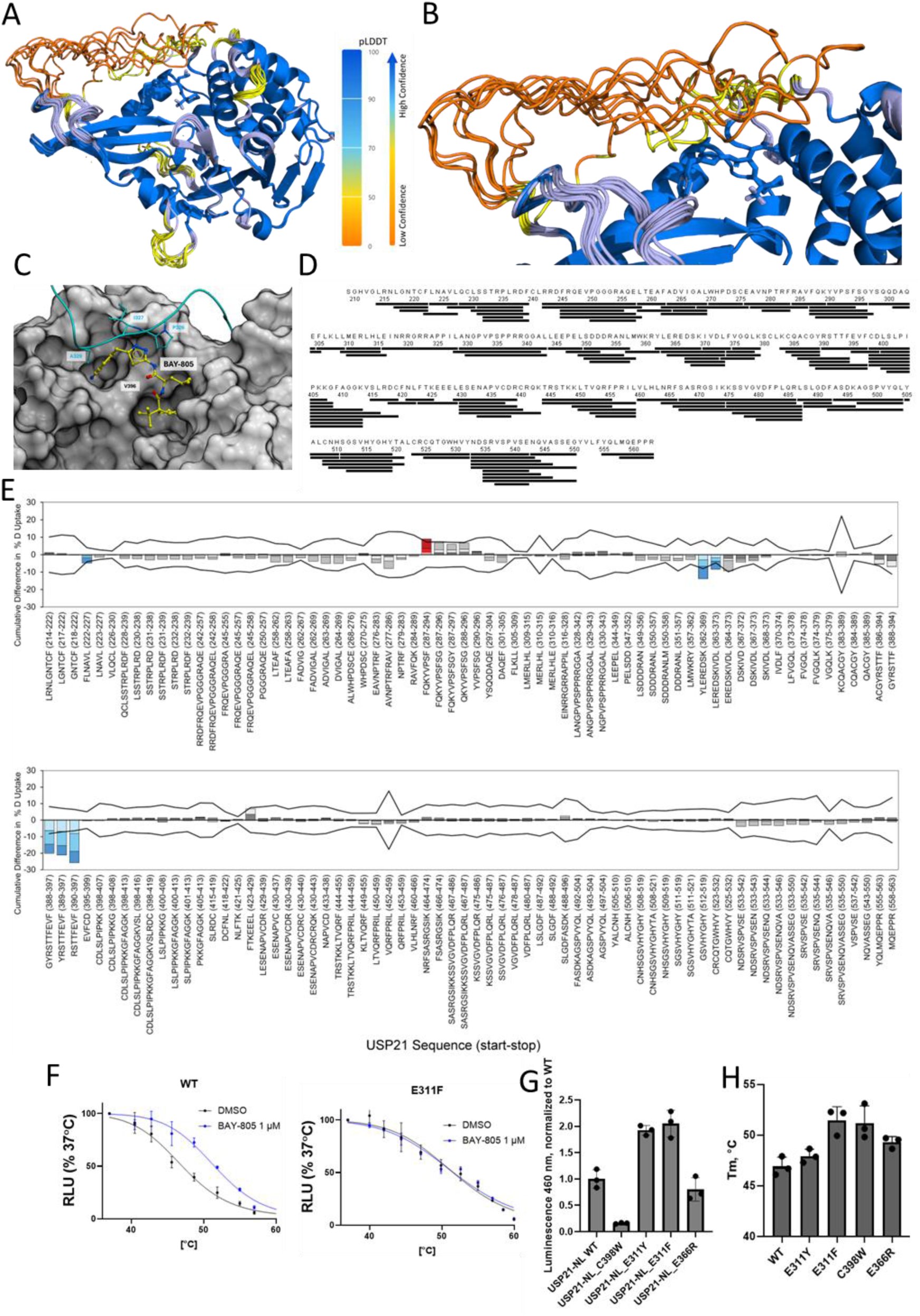
BAY-805 engages USP21 orthosteric site. (A) AF3 model ensemble of full-length USP21 in cartoon representation. pLDDT modeling confidence scores used to colour ensemble are indicated. (B) Closeup of the flexible loop region aa. 323 to 348 of USP21 shown in same format and colour scheme as A. (C) Surface representation of BAY-805 binding pocket with flexible loop shown as blue ribbon. Although the model suggested possible proximity of residues P326, I327, and A329 to the ligand, the uncertainty in this region limits definitive conclusions regarding direct contacts.. (D). Peptide Sequence Coverage of USP21. Using a 1:1 Nep2-Pep Affipro column, 136 USP21 peptides were generated, yielding a sequence coverage of 90.9% and average peptide-per-residue redundancy of 3.69. (E) Peptide-level, cumulative ΔHDX-MS. Labeling was performed at 0.25, 1, and 10 min. To be statistically significant, the stacked bar must exceed the cumulative propagated error at 3 sigma (line). Bars exceeding error are coloured blue (decreases) or red (increases), while insignificant bars are greyscale. (F) NalTSA testing of AF3 and HDX predicted residues that are required for BAY-805 binding. Thermal melting curves of NalTSA are shown for WT and E311F mutant of USP21. (G-H) Assessment of USP21 levels (G) and thermal stability (H) for mutants used in NalTSA. Luminescence intensities in E as proxy for nanoLuc tagged USP21 are shown.

**Figure S5.**
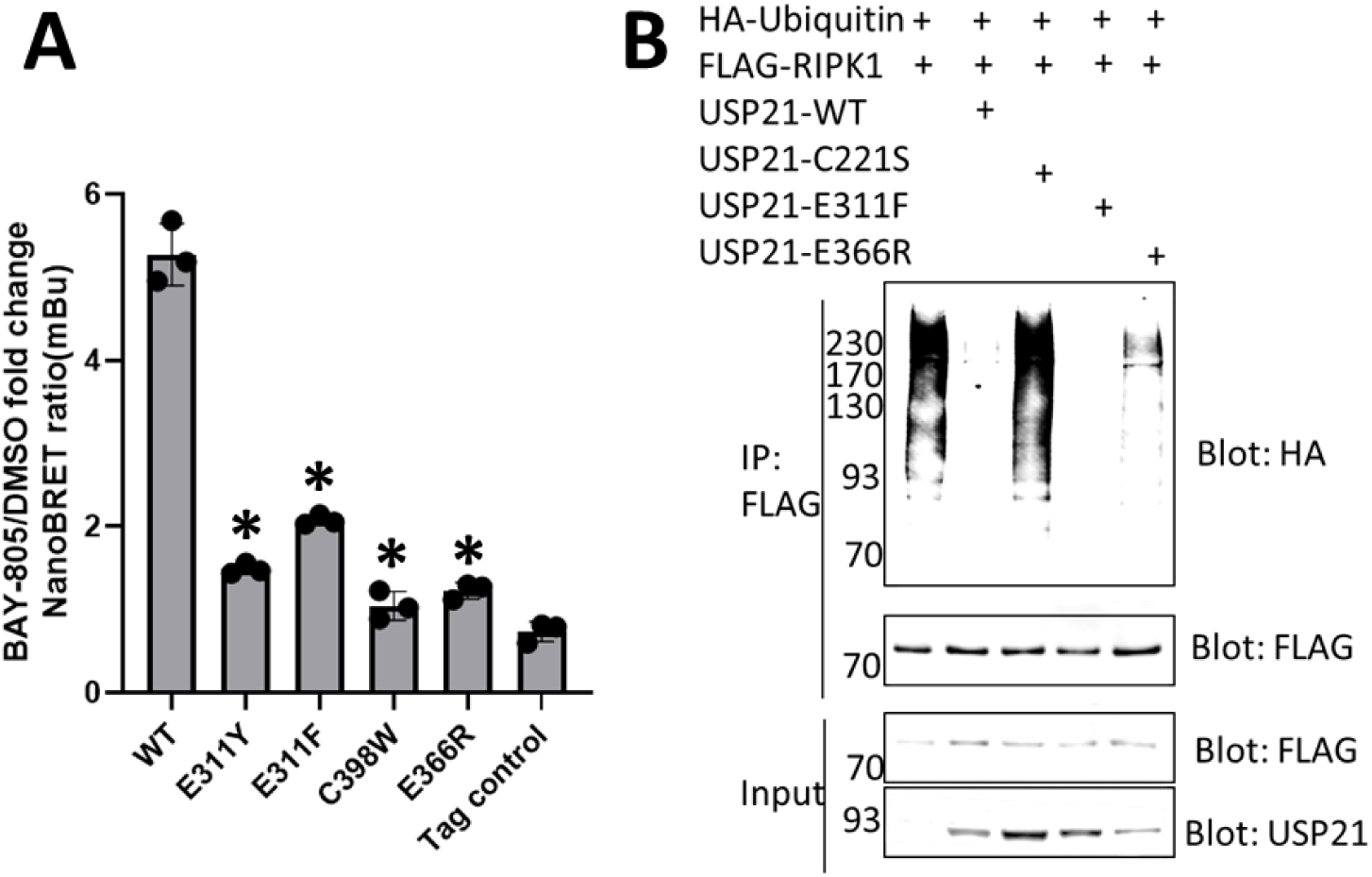
USP21 mutations that disrupt BAY-805 binding also disrupt BAY-805 induced USP21 association with KLHL12 irrespective of catalytic inhibition. (A) USP21 and KLHL12 interaction NanoBRET assay performed as in Fig 2D (B) USP21 E311F and E366R mutations do not abolish the catalytic activity. HEK293 cells were transfected with USP21 substrate Flag-RIPK1, HA tagged ubiquitin and various forms of USP21. Flag immunoprecipitates were blotted for HA ubiquitin. Experiment performed in 2 biological replicates, representative western blot is shown. C221S mutation of USP21 was used because C221A mutation resulted in very low protein levels.

**Figure S6. (Relating to Figure 6).**
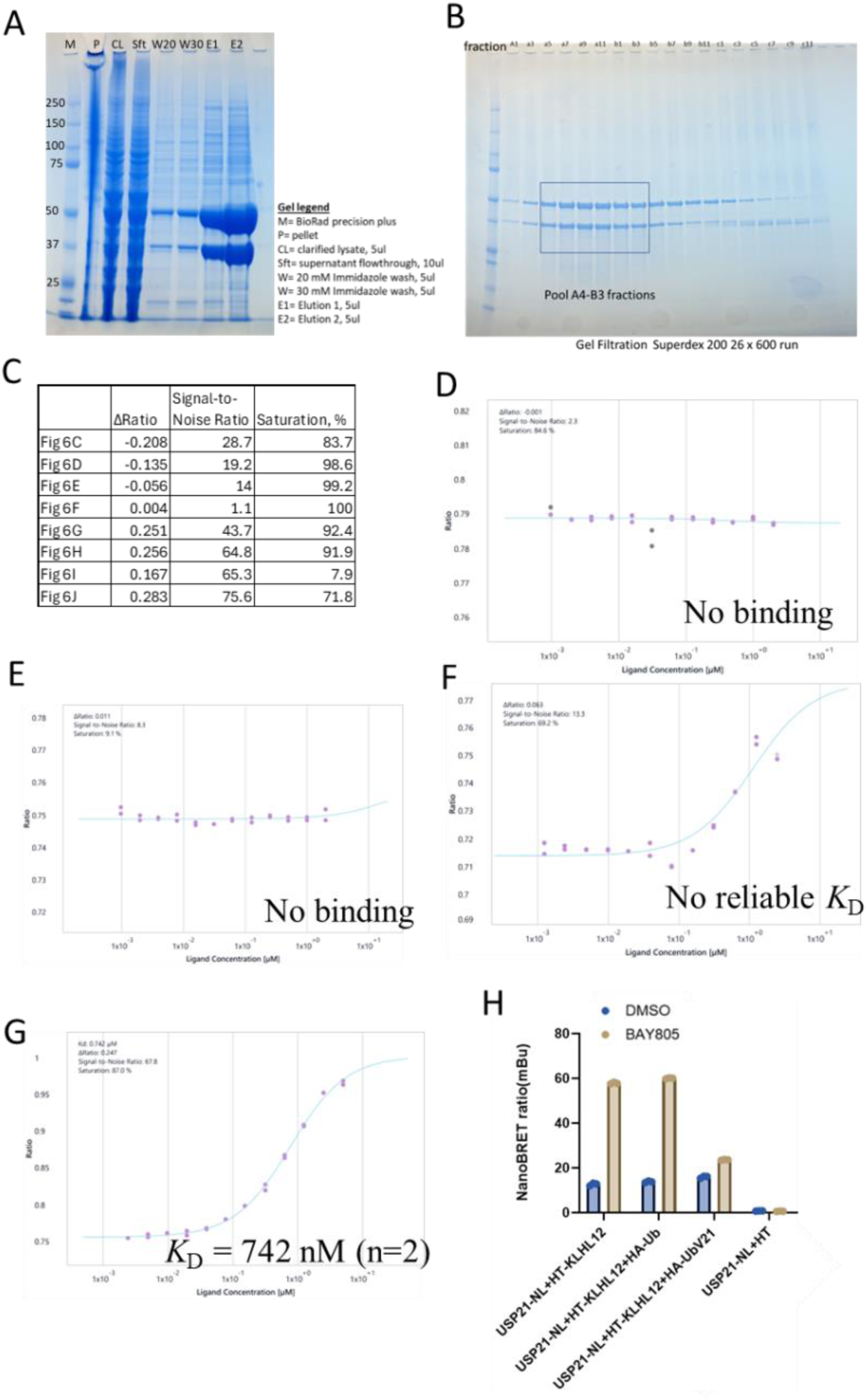
BAY-805 outcompetes ubiquitin and leads to *in vitro* USP21-KLHL12 binding. (A) N-terminally HiBit and LgBit tagged and His C-terminally tagged KELCH domain of KLHL12 protein expression and purification (protein used in Fig 6 G-J). Resulting two bands represent HiBit and LgBit tagged proteins. (B) Gel filtration of HiBit and LgBit His tagged Kelch domain of KLHL12 proteins. Coomassie staining of GF fractions. (C) Summary of SpS fitting parameters for data in Fig. 6C-J. (D-E) The N-terminally (D) or C-terminally (E) His-tagged KELCH domain of KLHL12 does not interact with BAY-805. Experiments were performed as in Fig. 6H. in the absence of USP21. (F) The N-terminally His-tagged HiBit-LgBit KELCH domain of KLHL12 does not interact with USP21 in the presence of ubiquitin. Experiments were performed as in Fig. 6I. (G) BAY-805 outcompetes ubiquitin and restores USP21 interaction with N-terminally His-tagged HiBit-LgBit KELCH domain of KLHL12. Experiments were performed as in Fig. 6J. (H) High affinity ubiquitin variant UbV21 prevents BAY-805 driven USP21 and KLHL12 association. NanoBRET was performed as in Fig 2D with/without wildtype ubiquitin or UbV21 co-transfections.

**Figure S7.**
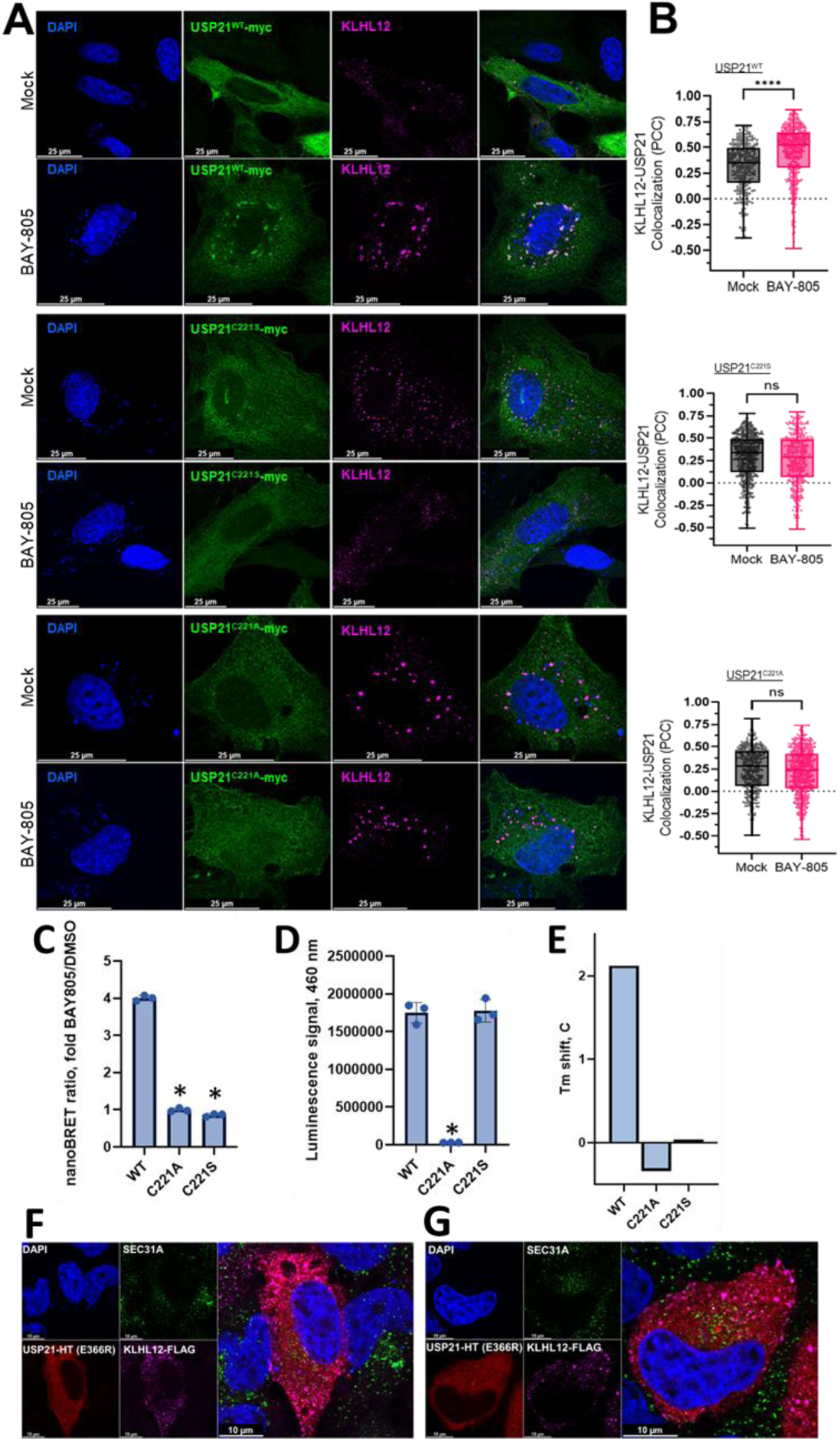
USP21 mutants that are deficient in BAY-805 induced KLHL12 association or catalytic activity are not stabilized by BAY-805 and they do not colocalize or interact with KLHL12. (A) Subcellular localization of USP21 catalytic cysteine mutations. USP21 WT or cysteine mutations (C221S, C221A) and KLHL12 were co-transfected into HeLa cells and subjected to BAY-805 treatment (1 μM, 30 min) in comparison to mock control. (B) Corresponding CellProfiler plots were generated from 12-16 cells per condition, analyzing the colocalization between USP21 and KLHL12 by Pearson’s Correlation Coefficient and. Asterisks (*) denote statistically significant findings for p<0.0001 (****), p<0.001 (***), p<0.05 (*); ns = not significant (C) Catalytic mutantions of USP21 abolish USP21-KLHL12 association in nanoBRET assay (D) C221A mutation results in lower expression levels of USP21. (E) USP21 catalytic mutants are not thermally stabilized by BAY-805. Experiment performed as in Figure 4H. (F) Subcellular localization of USP21 E366R. USP21 and KLHL12 were co-transfected into HeLa cells, mock control. (G) USP21 E366R does not colocalize with KLHL12 in respose to BAY-805. USP21 and KLHL12 were co-transfected into HeLa cells and subjected to BAY-805 treatment (1 μM, 30 min).

**Figure S8.**
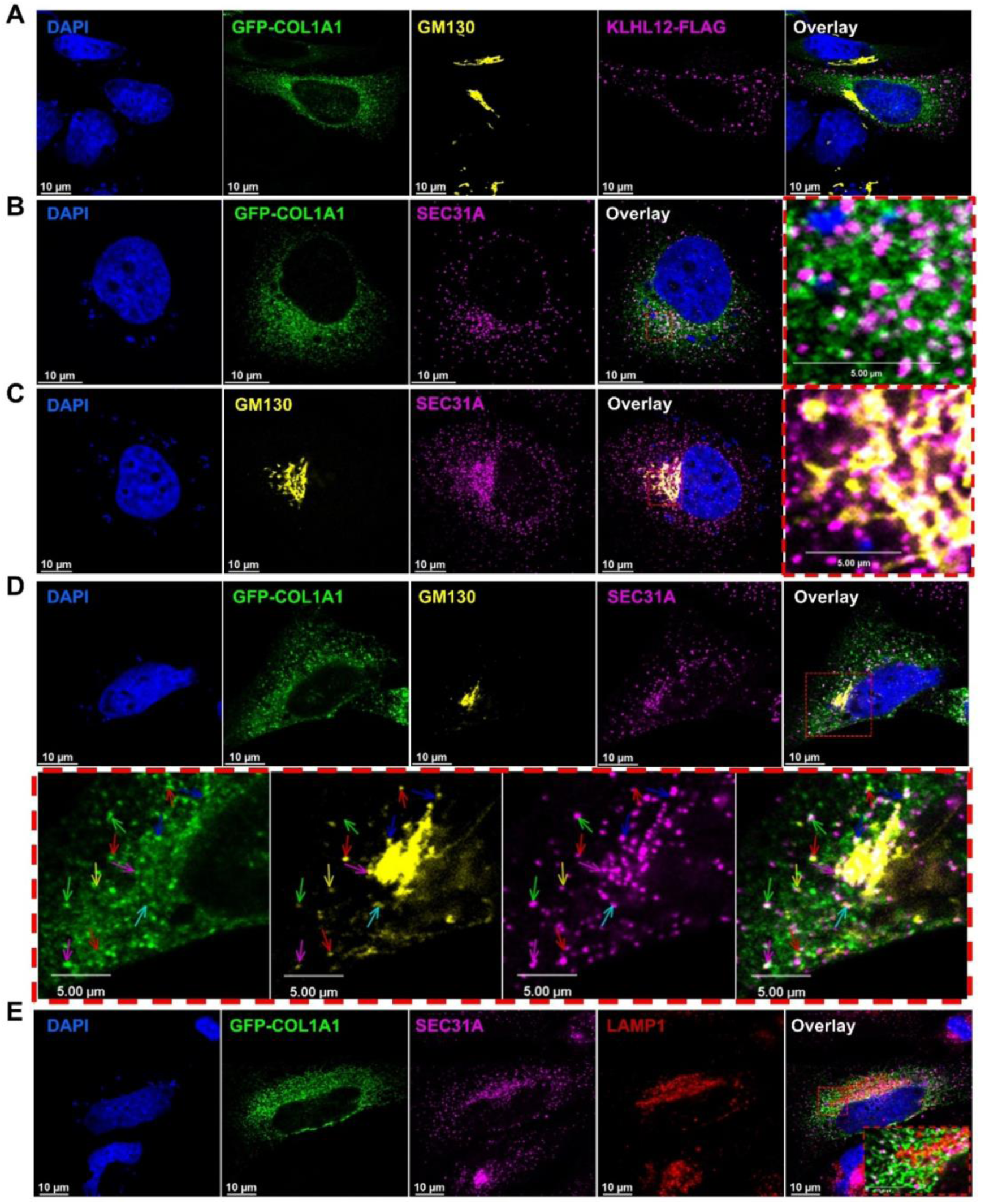
Subcellular localization of transiently overexpressed procollagen I (α1) in HeLa cells. Co-staining of endogenous markers for COPII vesicles (SEC31A), *cis*-Golgi network (GM130), and lysosomes (LAMP1) was carried out in combination with transfected GFP-COL1A1 with co-transfected KLHL12-FLAG. (A) Co-staining of GFP-COL1A1, *cis*-Golgi network (GM130), with co-transfected KLHL12-FLAG. (B) Co-staining of endogenous markers for COPII vesicles (SEC31A) with GFP-COL1A1. Zoomed-in regions are highlighted with a red dashed box (C) Co-staining control of endogenous markers for COPII vesicles (SEC31A) and *cis*-Golgi network (GM130). Zoomed-in regions are highlighted with a red dashed box (D) Co-staining of endogenous markers for COPII vesicles (SEC31A), *cis*-Golgi network (GM130) with GFP-COL1A1. Zoomed-in regions are highlighted with a red dashed box below panel D. (E) Co-staining of endogenous markers for COPII vesicles (SEC31A), lysosomes (LAMP1) with GFP-COL1A1. Zoomed-in regions are highlighted with a red dashed box

